# *In vivo*-directed evolution identifies AAV-WM04 as a next-generation vector for potent and durable hearing restoration in DFNB9

**DOI:** 10.64898/2026.03.11.710960

**Authors:** Yong Tao, Cenfeng Chu, Zhenzhe Cheng, Yilin Sun, Ying Chen, Huifang Zhang, Shuyue Bao, Boyu Yang, Baoyi Feng, Xianyu Huang, Yao Lu, Qiuxiang Yang, Xuegao Mao, Qifang Zhou, Chenxi Jin, Zhuo Duan, Guisheng Zhong, Hao Wu

## Abstract

Efficient and cell-specific gene delivery to cochlear inner hair cells (IHCs) remains a major challenge for inner ear gene therapy. Here, we identify and characterize a novel AAV2-derived capsid, AAV-WM04, that enables highly efficient and selective IHC transduction at low doses. Using an *in vivo*–directed evolution strategy, we generated a randomized AAV2 capsid library with 9–amino acid insertions and performed iterative selection in the adult mouse cochlea. Next-generation sequencing revealed enrichment of several variants, among which AAV-WM04 exhibited superior packaging efficiency and pronounced IHC tropism.

AAV-WM04 achieved near-complete IHC transduction throughout the cochlear axis in adult mice, outperforming clinically relevant vectors with minimal off-target expression and no detectable ototoxicity. Robust and exclusive IHC transduction was further validated in non-human primates following round window membrane delivery, underscoring translational potential.

Therapeutically, AAV-WM04 enabled efficient dual-AAV trans-splicing delivery of the large *OTOF* gene, resulting in uniform full-length otoferlin expression in IHCs. In a humanized *Otof* ^Q829X/Q829X^ mouse model, AAV-WM04 restored auditory function across a broad frequency range at relatively low doses and achieved durable hearing recovery.

Collectively, these findings establish AAV-WM04 as a next-generation IHC-targeted vector with high efficiency, safety, and cross-species applicability for precision gene therapy of hereditary hearing loss.

## Introduction

Autosomal recessive deafness type 9 (DFNB9) is a non-syndromic hereditary hearing loss caused by pathogenic variants in the *OTOF* gene, accounting for approximately 2%-8% of congenital deafness cases^1-5^. The encoded protein, otoferlin, functions as a calcium sensor predominantly expressed in inner hair cells (IHCs), where it mediates synaptic vesicle exocytosis and endocytosis, thereby enabling precise neurotransmission between IHCs and auditory nerve fibers^6-10^. Loss-of-function mutations in *OTOF* disrupt this process, impair synaptic transmission and ultimately lead to hearing loss.

Recent advances in adeno-associated virus (AAV)-mediated gene therapy have brought new hope to patients with *OTOF*-related deafness. Currently, several clinical trials, including ChiCTR2200063181, NCT05901480, NCT05821959, NCT05788536, NCT06370351, NCT06722170, and NCT06696456—are underway in China, the United States, and Europe to evaluate the safety and efficacy of *OTOF* gene therapy. Early clinical findings indicated significant improvements in the auditory capabilities of patients, supporting the potential of gene-based interventions for hereditary deafness and other forms of hearing disorders^11-14^. Nevertheless, a subset of patients fails to respond to therapy across clinical trials, highlighting the ongoing challenges in achieving consistent and durable hearing restoration through gene therapy.

To date, multiple AAV serotypes—including AAV1, AAV2quadY-F, AAV6, AAV8, AAV9, PHP.B, PHP.eB, and Anc80L65—have been used for *OTOF*-related gene therapy in mouse models^2,15-23^. In contrast, clinical trials have primarily adopted a dual-AAV recombination strategy, employing either AAV1 or Anc80L65 to accommodate the large size of the OTOF coding sequence^11-14^. Although Anc80L65 was applied clinically as a vector designed to enhance the transduction efficiency of IHCs, ectopic expression in outer hair cells has been observed in GFP transduction studies in non-human primates (NHPs)^19^. To mitigate off-target expression, previous studies have developed a hair cell-specific promoter, mMyo15, which significantly reduced ectopic expression in adult mice^17,19,24^.

For gene therapy, achieving sufficient expression levels and long-term persistence of therapeutic proteins is crucial, as these factors directly determine both the efficacy and durability of treatment. To attain adequate transgene expression, high vector doses are often required. However, elevated doses of AAV can provoke abnormal immune responses, exhibit poor tissue specificity, and compromise efficient transduction rates within target cells^25^. In animal models, AAV tends to accumulate in critical organs such as the liver and central nervous system in both mice and cynomolgus macaques, which poses risks of hepatotoxicity and neurotoxicity^26,27^. Therefore, the development of highly efficient and cell-targeted AAV vectors represents a promising strategy to achieve robust therapeutic efficacy at lower vector doses, thereby improving both safety and transduction efficiency in AAV-mediated gene therapy.

Therefore, achieving an optimal balance between therapeutic efficacy and safety remains a central challenge in gene therapy development. However, AAV vectors exhibiting genuine IHCs specificity are still unavailable, especially in adult animal models^28,29^. Engineering more efficient and tissue-selective delivery vectors thus represents a promising strategy to expand the therapeutic window and improve clinical translatability.

In this study, we employed an AAV peptide insertion library screening strategy^30^, utilizing the natural serotype AAV2 as the parental capsid and introducing randomized 9-amino acid peptides sequence, to identify a novel AAV variant, designated WM04, with selective tropism for IHCs. Considering that the human inner ear is fully developed *in utero* and the cochlea of P0-P3 mice is developmentally equivalent to the human cochlea prior to 26 weeks of gestational, we performed library screening in P30 mice, which corresponded to the late fetal or early postnatal stages of human inner ear maturation^22^.

WM04 achieved near-complete transduction of IHCs at substantially lower doses without the use of cell-type–specific promoters and exhibited minimal transduction of outer hair cells or supporting cells. In humanized *Otof* ^Q829X/Q829X^ mouse models, a single administration of WM04-OTOF resulted in dose-dependent restoration of auditory function, with both auditory brainstem response (ABR) thresholds and otoferlin expression remaining stable for up to eight weeks after treatment. Importantly, WM04 also demonstrated excellent safety and IHC specificity in non-human primates. In light of these preclinical findings, a clinical trial has been initiated to evaluate the safety, tolerability, and efficacy of WM04-OTOF in pediatric patients with hearing loss caused by *OTOF* mutations.

## Results

### A novel AAV capsid specifically targets IHCs following screening of AAV2-based capsid variants

Previous studies have demonstrated that AAV2 efficiently transduces cochlear IHCs, and several AAV2-based capsid mutants have been successfully applied in gene therapy for hereditary hearing loss. For example, the AAV2quadY-F mutant has been shown to restore hearing in *Otof* knockout mouse models. Building on these findings, we sought to engineer an AAV2-derived capsid with enhanced IHC specificity and transduction efficiency.

To this end, we generated an AAV2 capsid library by inserting random 9-amino acid peptides between amino acid positions N587 and R588 of the VP1 protein. The library was delivered into the cochleae of adult mice via posterior semicircular canal injection, followed by three iterative rounds of *in vivo* selection. AAV genomes recovered from cochlear tissues after each round (R1–R3) were analyzed by next-generation sequencing (NGS) (Figure 1A). Sequencing revealed progressive enrichment of the parental AAV2 sequence and four variants—AAV-201, AAV-203, AAV-205, and AAV-209—across all three selection rounds (Figure 1B). The relatively high abundance of AAV2 was likely attributable to residual plasmid contamination, consistent with its known cochlear tropism.

**Figure 1.**
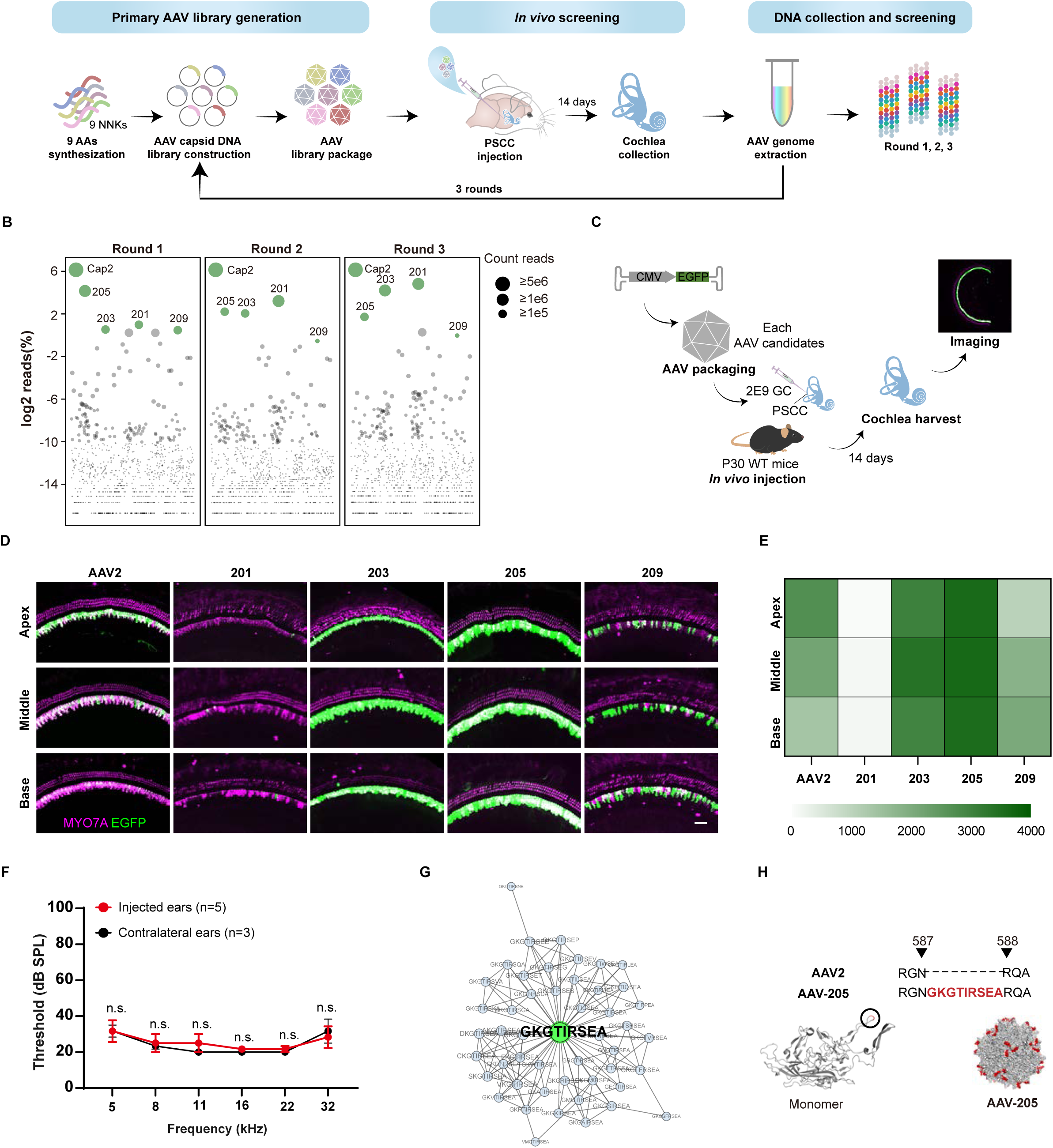
*In vivo* directed evolution screening identified AAV-WM04 as a potent and specific inner hair cell-targeted vector. Schematic of *in vivo*–directed evolution process used to screen cochlear enriched variants. The library, created by inserting a random nine-amino acid peptide between position 587 and 588 position of AAV2 capsid protein, was packaged and injected into the cochleae through PSCC. 14 days post-injection, cochleae were collected to extract the genomic DNA. Next generation DNA sequencing of the genomic DNA was performed to analyze the enrichment of peptide motifs, and PCR-amplified from genomic DNA was performed to enrich viral cap genes (representing the AAV variants from the library that successfully transduced cochleae) for cloning and repackaging. Three rounds of *in vivo* selections were carried out. **B.** Distribution of variants recycled from each round weas shown with capsid libraries sorted by descending order of the NGS reads (log2 scale). The top 5 reads—AAV2, 201, 203, 205, and 209—were mapped as green dots on the plot. **C.** Schematic illustration of the experimental design. GFP-expressing AAV2, AAV-201, AAV-203, AAV-205, and AAV-209 were independently packaged and delivered into the cochleae of C57BL/6J mice via PSCC injection at P30. Cochleae were harvested 14 days post-injection and subjected to immunostaining. **D.** Representative confocal images of cochleae injected with AAV candidates in adult mice. AAV capsid candidates were packaged with CMV-eGFP and verified *in vivo* in adult mice. Each AAV was injected into the mouse cochlea via PSCC at the same dose (5x10^9^ GCs per ear), and cochlear tissue was collected 14 days later. Green: EGFP protein; magenta: MYO7A. Scale bar: 20 μm. **E.** Relative fluorescence intensity of EGFP-positive IHCs in the apical, middle and basal region of cocleae corresponding to **C**. Data were shown as mean. **F.** ABR thresholds of adult WT mice at 14 days post-injection with AAV-WM04-CMV-EGFP (2x10^9^ GCs per ear). Data were shown as mean ± SEM. Significance tests were performed between AAV-WM04 and other AAV serotypes. *P* Value is calculated by Student’s *t*-test. n.s. means no significant difference. **G.** Clustering analysis of 205 family corresponding to the top 4 peptides from positively enriched variants across all three rounds. **H.** Molecular model of AAV-WM04 containing the insertion peptide GKGTIRSEA (shown in red) after amino acid 587 of AAV2, monomeric form was on left and 60-meric form was on right.

To evaluate the transduction properties of the enriched variants, each capsid was packaged with a CMV-EGFP reporter and injected into the cochleae of postnatal day 30 (P30) C57BL/6J wild-type mice at an equal dose of 2 × 10^9^ genome copies (GC) per ear. Cochlear tissues were harvested two weeks after injection for EGFP analysis (Figure 1C). At this dose, both AAV-203 and AAV-205 achieved near-complete IHC transduction throughout the apical, middle, and basal turns, reaching 100% efficiency. In contrast, AAV-201 exhibited minimal transduction (3.70%, 4.27%, and 4.28%), while AAV-209 showed moderate efficiency (49.44%, 59.82%, and 74.23%) across the respective cochlear regions (Figure 1D; Figure S1A).

Although AAV-203 and AAV-205 displayed comparable IHC transduction rates, quantitative analysis revealed that AAV-205 drove significantly stronger and more uniform EGFP expression across the cochlear axis. The relative fluorescence intensities of EGFP mediated by AAV-205 were 3,597.87 ± 73.61 (apex), 3,624.52 ± 25.85 (middle), and 3,560.66 ± 148.33 (base), whereas AAV-203 yielded lower intensities of 3,041.10 ± 141.57, 3,396.07 ± 89.87, and 2,976.41 ± 324.78, respectively (Figure 1E).

Assessment of viral packaging efficiency further demonstrated that AAV-205 achieved the highest production titer (1.04 × 10^13^ GC/mL), exceeding those of AAV-201 (8.09 × 10^12^ GC/mL), AAV-203 (2.90 × 10^12^ GC/mL), and AAV-209 (7.10 × 10^11^ GC/mL) (Figure S1B). Importantly, AAV-205 exhibited a favorable safety profile in adult mice, as ABR thresholds remained unchanged following inner ear injection at 1.04 × 10^13^ GC/mL (Figure 1F). Consistent with its localized cochlear transduction, no EGFP signal was detected in the brain or liver, indicating minimal systemic dissemination (Figure S2).

To further investigate the sequence relationships among enriched variants, reverse Hamming distances—defined as the number of shared amino acids between two peptide insertions—were calculated for all unique variants detected across the R1–R3 selection rounds. Network clustering based on these distances revealed four dominant capsid families, comprising 57, 49, 46, and 26 variants, respectively (Figure S3; Figure 1G). Notably, AAV-201 (57 variants), AAV-203 (49 variants), AAV-205 (46 variants), and AAV-209 (26 variants) occupied central hub positions within four distinct clusters, representing the major evolutionary lineages selected during *in vivo* screening.

AAV-205 contains a 9-amino acid insertion (GKGTIRSEA) at the engineered capsid site (Figure 1H). Structural modeling suggested that this insertion alters the conformation of loop 4, potentially disrupting key arginine residues involved in AAV2 primary receptor binding and thereby modulating viral tropism. AAV-WM04 retained canonical VP1/VP2/VP3 stoichiometry (Figure S4) while exhibiting enhanced IHC transduction efficiency and high vector yield. Based on its superior transduction efficiency, expression strength, and packaging yield, AAV-205 was selected for further characterization and designated AAV-WM04.

Collectively, these results identify AAV-WM04 as a robust and safe IHC-targeted vector suitable for further development in next-generation gene therapy for hereditary hearing loss.

### Intra-cochlea delivery of AAV-WM04 enables robust and low-dose transduction of IHCs in mice

To further evaluate the transduction efficiency of AAV-WM04 in IHCs, we compared its performance with two clinically relevant AAV serotypes, AAV1 and Anc80L65, across a range of vector doses. At a dose of 2 × 10^9^ GCs per ear, AAV-WM04 transduced nearly all IHCs throughout the cochlea, whereas AAV1 and Anc80L65 achieved transduction efficiencies of approximately 60% across cochlear regions (Figure S5A, B).

Strikingly, at a lower dose of 5 × 10^8^ GCs per ear (n = 3), AAV-WM04 maintained complete (100%) IHC transduction across apical, middle, and basal turns, whereas both AAV1 and Anc80L65 exhibited a marked decline in transduction efficiency (Figure 2A, B). Specifically, AAV1 achieved transduction efficiencies of 80.93 ± 0.40%, 77.50 ± 1.16%, and 55.07 ± 7.28% in the apical, middle, and basal turns, respectively, while Anc80L65 showed substantially lower efficiencies of 31.10 ± 23.04%, 36.40 ± 18.36%, and 35.00 ± 19.54% (Figure 2B). Notably, even at a further reduced dose of 1 × 10^8^ GCs per ear (n = 3), AAV-WM04 sustained near-complete transduction of IHCs across the entire cochlear axis.

**Figure 2.**
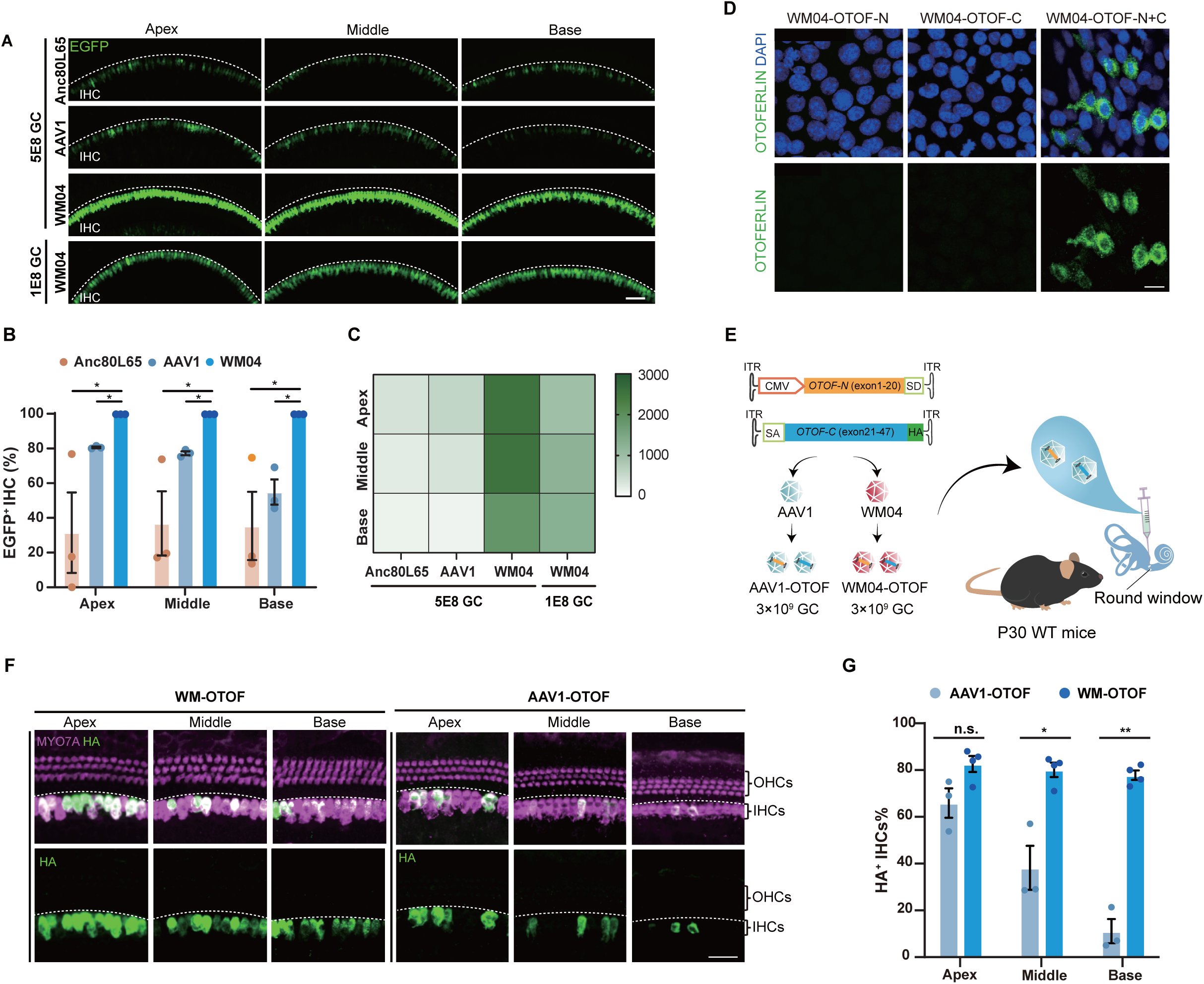
AAV-WM04 exhibits superior inner hair cell transduction efficiency at low doses. **A.** Representative confocal images of cochleae from P30 wild-type (WT) mice injected with AAV-WM04-CMV-EGFP (5 × 10^8^ or 1 × 10^8^ GCs), Anc80L65-CMV-EGFP (5 × 10^8^ GCs), or AAV1-CMV-EGFP (5 × 10^8^ GCs). EGFP expression is shown in green. IHCs, inner hair cells. Scale bar, 20 μm. **B.** Quantification of the percentage of EGFP-positive IHCs in apical, middle, and basal cochlear turns corresponding to **A**. Data are presented as mean ± SEM (n = 3 mice per group). Statistical comparisons were performed between AAV-WM04 and other AAV serotypes using Student’s *t*-test. n.s., not significant; *P* < 0.05. **C.** Relative fluorescence intensity of EGFP-positive IHCs in the apical, middle, and basal regions corresponding to **A**. Data are shown as mean values. **D.** Representative confocal images of HEK293 cells infected with WM04-OTOF-N alone, WM04-OTOF-C alone, or a combination of WM04-OTOF-N and WM04-OTOF-C. Otoferlin expression is shown in green; nuclei are labeled with DAPI (blue). Scale bar, 10 μm. **E.** Schematic illustration of the experimental design for dual-AAV OTOF delivery. Recombinant AAV-OTOF-N and AAV-OTOF-C-HA vectors were independently packaged and co-delivered into the mouse cochlea via RWM administration. **F.** Representative confocal images of cochleae from WT mice 14 days after injection with AAV-WM04-OTOF-HA or AAV1-OTOF-HA (3 × 10^9^ GCs). HA-tagged OTOF expression is shown in green; MYO7A is shown in magenta. OHCs, outer hair cells; IHCs, inner hair cells. Scale bar, 20 μm. **G.** Quantification of the percentage of HA-positive IHCs in apical, middle, and basal cochlear regions corresponding to **F**. Data are presented as mean ± SEM (n = 3–4 mice). Statistical significance was determined using Student’s *t*-test. n.s., not significant; *P* < 0.05; *P* < 0.01.

Consistent with its superior transduction efficiency, AAV-WM04 also drove markedly stronger transgene expression than AAV1 and Anc80L65. Quantitative analysis of EGFP fluorescence intensity revealed that AAV-WM04 mediated robust expression in the apical, middle, and basal turns (2,512.88 ± 97.49, 2,538.45 ± 34.48, and 1,854.72 ± 24.64, respectively). In contrast, AAV1 yielded lower fluorescence intensities (571.40 ± 45.26, 362.68 ± 34.53, and 128.12 ± 14.77), while Anc80L65 produced weaker and less uniform expression across cochlear regions (364.55 ± 22.23, 224.20 ± 14.91, and 141.62 ± 21.84) (Figure 2C).

Taken together, these results demonstrate that AAV-WM04 combines highly-efficient transduction and robust transgene expression in IHCs even at substantially reduced doses, outperforming current clinically relevant AAV serotypes. AAV-WM04 can be a promising platform for safe and effective inner ear gene therapy.

### Dual AAV-WM04-mediated delivery of *OTOF* enables efficient recombination in IHCs

To evaluate the therapeutic potential of AAV-WM04 for IHC-targeted gene therapy, we examined its performance in the context of *OTOF* gene transfer. The human *OTOF* coding sequence spans 5,991 bp, exceeding the packaging capacity of a single AAV vector (∼4.7 kb). Therefore, consistent with previous reports, we employed a dual-AAV trans-splicing strategy targeting exons 20 and 21 of *OTOF* (Figure S6A).

To validate trans-splicing and recombination efficiency, pAAV-CMV-OTOF-N and pAAV-OTOF-C-HA were packaged into AAV-WM04 and mixed at a 1:1 ratio (designated WM04-OTOF-HA). Western blotting and immunofluorescence analyses confirmed robust expression of full-length otoferlin in HEK293 cells infected with WM04-OTOF-HA, indicating efficient vector recombination and protein reconstitution (Figure 2D).

We next compared *in vivo OTOF* expression mediated by AAV-WM04 and the clinically relevant AAV1 serotype. Equal viral doses (3 × 10^9^ GCs per ear) of AAV-WM04-OTOF-HA or AAV1-OTOF-HA were delivered into the cochleae of adult wild-type mice via PSCC injection (Figure 2E). Fourteen days post-injection, HA-tagged full-length OTOF expression was detected exclusively in IHCs following AAV-WM04 administration and at substantially higher recombination efficiency than that achieved by AAV1 (Figure 2F). Importantly, this high recombination efficiency was achieved at a relatively low viral dose, underscoring the superior transduction potency of AAV-WM04 and suggesting a reduced vector burden for therapeutic application.

Quantitative analysis revealed that WM04-OTOF-HA achieved fusion efficiencies of 82.60 ± 3.40%, 80.15 ± 3.10%, and 77.83 ± 2.07% in the apical, middle, and basal turns, respectively. In contrast, AAV1-mediated recombination efficiencies were 65.93 ± 6.31%, 38.20 ± 9.40%, and 11.13 ± 5.17% across the corresponding regions (Figure 2G). Notably, AAV1 exhibited a progressive decline in recombination efficiency from apex to base, consistent with the limited high-frequency hearing recovery reported in previous studies. A similar gradient has been reported for Anc80L65-mediated *OTOF* delivery. In contrast, AAV-WM04 supported uniform recombination across the cochlear axis, particularly in the high frequency region, providing a mechanistic basis for the robust recovery of auditory function in subsequent functional analyses.

Collectively, these results demonstrate that dual AAV-WM04 enables highly efficient and uniform OTOF recombination in IHCs at low vector doses, establishing a robust molecular foundation for durable and broadband hearing restoration.

### AAV-WM04 restores hearing in *Otof* ^Q829X/Q829X^ mice at low doses

To evaluate therapeutic efficacy in a deaf animal model, we used a humanized *Otof*^Q829X/Q829X^ mouse model, in which exon 21 of *OTOF* was deleted using CRISPR-Cas9 guided by four sgRNAs flanking exon 21 and replaced with the corresponding human exon containing the Q829X mutation^31^.The humanized *Otof* ^Q829X/Q829X^ mouse model, which display profound deafness by 4 weeks of age with preserved outer hair cell function, thereby recapitulating the human *OTOF*-deficiency phenotype.

Equal doses of dual-AAV vectors (total 5 × 10^9^ GCs per ear, comprising 2.5 × 10^9^ GCs of AAV-OTOF-N and 2.5 × 10^9^ GCs of AAV-OTOF-C) packaged in either AAV2/1 (AAV1-OTOF) or AAV-WM04 (AAV-WM04-OTOF) capsids were delivered via round-window membrane (RWM) injection, a clinically relevant delivery route (Fig. 3A), in P30 *Otof* ^Q829X/Q829X^ homozygous mice (homo mice). ABRs to click or tone bursts stimuli were subsequently recorded to assess therapeutic efficacy.

**Figure 3.**
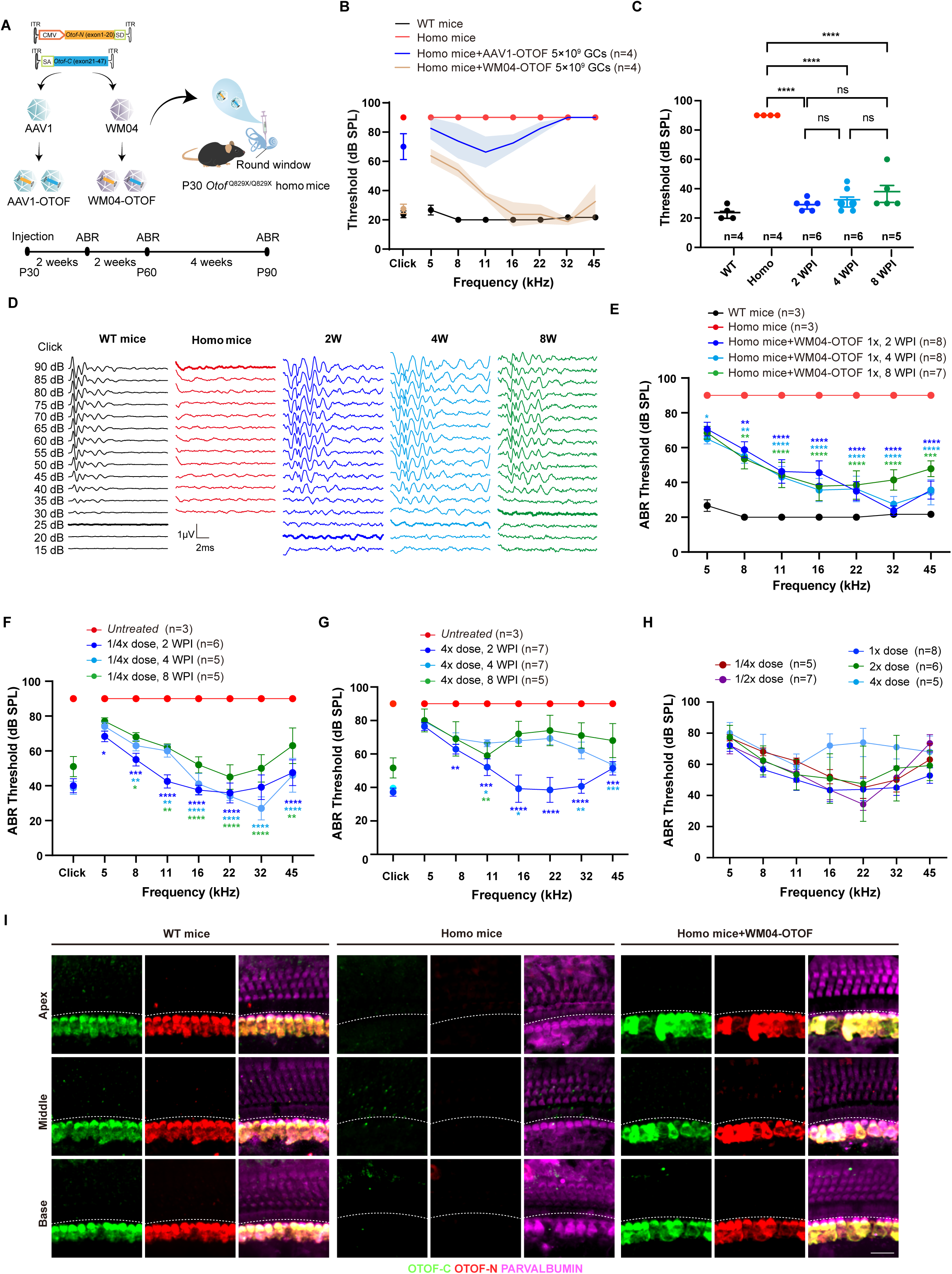
WM04-*OTOF* injection restored auditory function in P30 *Otof* ^Q829X/Q829X^ mice. **A.** Schematic overview of the pharmacodynamic study design. Recombinant dual-AAV vectors encoding OTOF-N and OTOF-C were independently packaged into the AAV-WM04 capsid and delivered into the cochleae of P30 *Otof* ^Q829X/Q829X^ homozygous mice via round window membrane injection. WM04-OTOF was administered at doses of 1.25 × 10^9^, 2.5 × 10^9^, 5 × 10^9^, 1 × 10^10^, and 5 × 10^10^ GCs per ear. AAV1-packaged OTOF (AAV1-OTOF, 5 × 10^9^ GCs) served as a control. **B.** Tone-burst ABR thresholds of P30 wild-type mice, untreated homozygous mice, and homozygous mice treated with WM04-OTOF (5 × 10^9^ or 1 × 10^10^ GCs) or AAV1-OTOF (5 × 10^9^ GCs) at 2 weeks post-injection (n = 4 per group). Data are presented as mean ± SEM. **C.** Click-evoked ABR thresholds in P30 wild-type mice (black, n = 4), untreated homozygous mice (red, n = 4), and homozygous mice treated with 5 × 10^9^ GCs of WM04-OTOF at 2 weeks (blue, n = 4), 4 weeks (light blue, n = 4), and 8 weeks (green, n = 4) post-injection. Data are shown as mean ± SEM. **D.** Representative click-evoked ABR waveforms from P30 wild-type mice (black), untreated homozygous mice (red), and homozygous mice treated with 5 × 10^9^ GCs of WM04-OTOF at 2 weeks (blue, n = 6), 4 weeks (light blue, n = 6), and 8 weeks (green, n = 5) post-injection. Bold traces indicate ABR thresholds. **E.** Frequency-specific tone-burst ABR thresholds of P30 wild-type mice (black, n = 3), untreated homozygous mice (red, n = 4), and homozygous mice treated with 5 × 10^9^ GCs of WM04-OTOF at 2 weeks (blue, n = 8), 4 weeks (light blue, n = 8), and 8 weeks (green, n = 7) post-injection. Data are shown as mean ± SEM. Statistical significance was determined by two-way ANOVA compared with untreated homozygous mice. *, **, ***, and **** indicate *P* < 0.05, 0.01, 0.001, and 0.0001, respectively. **F.** Click and tone-burst ABR thresholds of untreated homozygous mice (red, n = 4) and homozygous mice treated with a low dose of WM04-OTOF (1.25 × 10^9^ GCs, 1/4×) at 2 weeks (blue, n = 6), 4 weeks (light blue, n = 5), and 8 weeks (green, n = 5) post-injection. Data are shown as mean ± SEM. Statistical significance was assessed by two-way ANOVA compared with untreated homozygous mice. **G.** Click and tone-burst ABR thresholds of untreated homozygous mice (red, n = 4) and homozygous mice treated with a high dose of WM04-OTOF (2 × 10^10^ GCs, 4×) at 2 weeks (blue, n = 7), 4 weeks (light blue, n = 7), and 8 weeks (green, n = 5) post-injection. Data are shown as mean ± SEM. Two-way ANOVA compared with untreated homozygous mice. **H.** Dose–response analysis of tone-burst ABR thresholds at 4 weeks post-injection in homozygous mice treated with WM04-OTOF at 1.25 × 10^9^ (1/4×, n = 5), 2.5 × 10^9^ (1/2×, n = 7), 5 × 10^9^ (1×, n = 8), 1 × 10^10^ (2×, n = 6), and 2 × 10^10^ GCs (4×, n = 5). Data are shown as mean ± SEM. **I.** Representative confocal images of apical, middle, and basal cochlear turns from P30 wild-type mice, untreated homozygous mice, and homozygous mice treated with 5 × 10^9^ GCs of WM04-OTOF. Green, OTOF-N; red, OTOF-C; magenta, parvalbumin. Scale bar, 20 μm.

Two weeks after treatment, click-evoked ABR thresholds in AAV-WM04-OTOF-treated ears were restored to 27.5 ± 6.5 dB, closely approaching wild-type levels (23.8 ± 4.8 dB) and were markedly improved compared with AAV1-OTOF-treated ears (70.0 ± 8.9 dB). Frequency-specific ABR analysis showed that AAV1-OTOF partially restored hearing at low to mid frequencies (5–22 kHz) but failed to rescue high-frequency hearing (32–45 kHz). In contrast, AAV-WM04-OTOF achieved robust hearing recovery across a substantially broader frequency range, restoring thresholds to near wild-type levels from 16 to 45 kHz (Figure 3B).

To assess the durability of hearing restoration, ABRs were recorded at multiple time points following administration of the 1× dose (2.5 × 10^9^ GCs each of AAV-WM04-OTOF-N and AAV-WM04-OTOF-C). Click-evoked ABR thresholds were 32.5 ± 8.2 dB and 38.5 ± 13.0 dB at 4 and 8 weeks post-injection (WPI), respectively, with no significant differences relative to 2 WPI (Figure 3C). ABR waveforms further confirmed sustained auditory function 8 WPI (Figure 3D).

Quantitative analysis revealed stable threshold recovery from 2 to 8 WPI across most frequencies, except at 5 kHz. At low frequencies (5–11 kHz), thresholds ranged from 46.5 to 70.5 dB. At mid frequencies (11–22 kHz), thresholds ranged from 33.5–46.5 dB, 38.0–48.0 dB, and 43.9–50.0 dB at 2, 4, and 8 WPI, respectively. Notably, at high frequencies (32 and 45 kHz), thresholds at 2 WPI (22.0 and 31.5 dB) were comparable to wild-type values (21.7 and 21.7 dB) (Figure 3E).

Dose-response studies demonstrated that lower doses—1/2× (1.25 × 10^9^ GCs each of AAV-OTOF-N and -C) and 1/4× (6.25 × 10^8^ GCs each)—still mediated substantial hearing restoration, although high-frequency recovery was modestly reduced compared with the 1× dose (Figure 3F; Figure S7A). In contrast, higher doses (2× and 4×) did not further enhance efficacy and were associated with mild, time-dependent threshold elevations at high frequencies (Figure 3G; Figure S7B), indicating a plateau in therapeutic benefit.

Comparison of different doses at 4 weeks post-injection revealed comparable ABR thresholds across the 1/4×, 1/2×, 1×, and 2× dose groups. In contrast, although the highest tested dose (4×) did not affect auditory function at 2 WPI, it was associated with elevated ABR thresholds at middle frequencies at later time points (Figure 3H; Figure S8), suggesting a reduced safety margin at supratherapeutic exposure.

Consistent with functional recovery, otoferlin protein was undetectable in untreated *Otof* ^Q829X/Q829X^ mice but was robustly re-expressed throughout the cochlea 8 weeks after AAV-WM04-OTOF administration. Immunofluorescence analysis confirmed efficient re-expression of both OTOF-N and OTOF-C in IHCs, with full-length otoferlin detected in up to 100% of virus-infected IHCs (Figure 3I).

Collectively, these results demonstrate that AAV-WM04 enables efficient, durable, and broadband hearing restoration in humanized *Otof* ^Q829X/Q829X^ mice at low vector doses. Compared with the clinically used AAV1 serotype, AAV-WM04 achieves superior high-frequency rescue, maintains stable auditory function for at least eight weeks, and exhibits a favorable dose–response profile with a defined therapeutic window. These findings establish AAV-WM04 as a highly potent and translationally promising vector for *OTOF* gene therapy.

### Safety and biodistribution assessment of dual AAV-WM04-OTOF following intracochlear delivery

To evaluate the safety profile and biodistribution of dual AAV-WM04-OTOF vectors following a single intracochlear administration, we performed a comprehensive toxicity and distribution study in mice. Dual AAV-WM04-OTOF vectors were delivered into the inner ears of P30 C57BL/6J wild-type mice via RWM approach. Auditory function, motor coordination, learning and memory, as well as histopathological changes were assessed at 2 and 4 weeks post-injection (Figure 4A).

**Figure 4.**
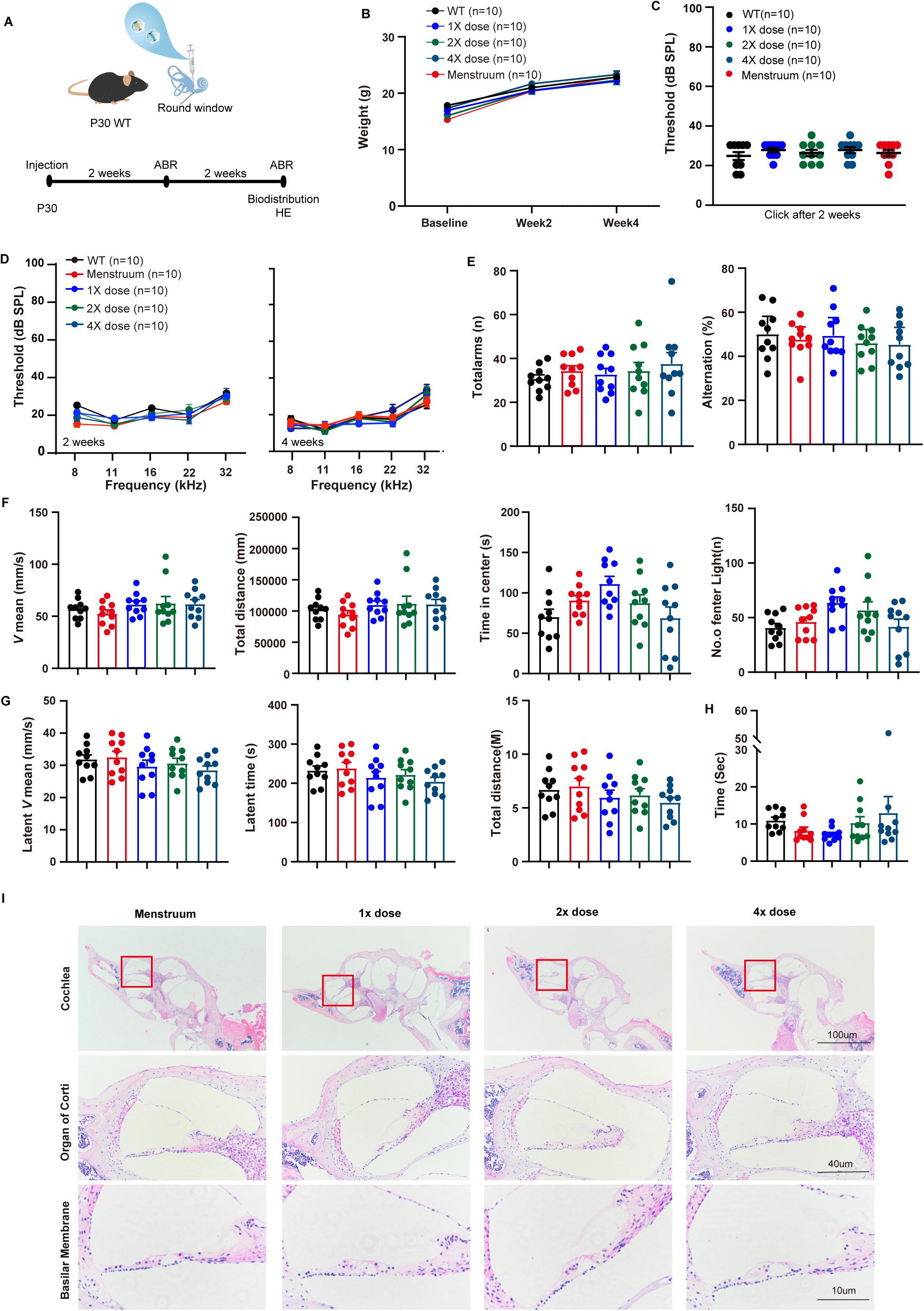
Safety evaluation of dual AAV-WM04-CMV-OTOF in adult wild-type mice. **A.** Schematic overview of the safety assessment of dual AAV-WM04-CMV-OTOF in P30 WT mice. Dual AAV-WM04-OTOF vectors were delivered via RWM injection. ABR were assessed at 2 and 4 weeks post-injection, while biodistribution analysis and H&E staining were performed at 4 weeks post-injection. **B.** Body weight measurements of adult WT mice and WT mice injected with menstruum or different doses of dual AAV-WM04-CMV-OTOF. Data are shown as mean ± SEM. **C.** Click-evoked ABR thresholds of adult WT mice (black, n = 10) and WT mice injected with menstruum (red, n = 10), 1× dose (blue, n = 10), 2× dose (green, n = 10), or 4× dose (sky blue, n = 10) of dual AAV-WM04-CMV-OTOF at 2 weeks post-injection. **D.** Tone-burst ABR thresholds measured at 2 and 4 weeks after injection of different doses of dual AAV-WM04-CMV-OTOF. Data are presented as mean ± SEM. **E.** Y-maze test assessing total arm entries and spontaneous alternation behavior in WT mice and WT mice injected with menstruum or different doses of dual AAV-WM04-CMV-OTOF. Data are shown as mean ± SEM. **F.** Open-field test evaluating mean velocity, total distance traveled, time spent in the center, and number of entries into the light zone in WT mice and WT mice injected with menstruum or different doses of dual AAV-WM04-CMV-OTOF. Data are shown as mean ± SEM. **G.** Rotarod performance assessing latency to fall, mean velocity, and total distance traveled in WT mice and WT mice injected with menstruum or different doses of dual AAV-WM04-CMV-OTOF. Data are shown as mean ± SEM. **H.** Pole test assessing motor coordination in WT mice and WT mice injected with menstruum or different doses of dual AAV-WM04-CMV-OTOF. Data are shown as mean ± SEM. **I.** Representative HE-stained sections of the inner ear 2 weeks after injection of menstruum or dual AAV-WM04-OTOF at 1×, 2×, or 4× doses. The organ of Corti and basilar membrane are shown from the apical turn (red boxed region). Scale bars are indicated.

To assess whether dual AAV-WM04-OTOF administration affected general health, body weight was monitored in mice receiving vehicle control (menstruum, defined as the formulation buffer used to dissolve AAV-WM04-OTOF-N and AAV-WM04-OTOF-C), a therapeutic dose (1×; total 5 × 10^9^ GCs), or supratherapeutic doses (2× and 4×) at P30. To ensure data consistency, body weights of animals within each sex and treatment group were maintained within ± 20% of the group mean. No significant differences in body weight were observed among the menstruum, 1×, 2×, and 4× dose groups throughout the observation period (Figure 4B).

ABR testing performed two weeks after injection revealed no significant changes in click-evoked ABR thresholds across all treatment groups compared with uninjected wild-type controls (Figure 4C), indicating that neither the surgical procedure nor dual AAV-WM04-OTOF delivery impaired baseline auditory function. Similarly, frequency-specific ABR thresholds showed no significant differences among groups at most tested frequencies (Figure 4D), representing a non-adverse, dose-limiting observation that did not affect broadband auditory function.

To assess potential off-target neurological effects, motor coordination and cognitive function were evaluated four weeks post-injection using the rotarod test, pole test, Y-maze test, and open field test. No significant differences were observed among the menstruum, 1×, 2×, 4×, and untreated wild-type groups in locomotor activity, exploratory behavior, learning, or memory (Figure 4E–H).

Histopathological analysis was conducted four weeks after surgery to examine potential tissue damage. Hematoxylin and eosin (H&E) staining of cochlear sections revealed preserved inner ear morphology and intact organ of Corti architecture across all treatment groups, with no evidence of hair cell loss or structural abnormalities (Figure 4I). Given the potential for systemic dissemination of AAV vectors, major organs were also examined. Microscopic analysis of H&E-stained sections from the brain, lymph node, liver, kidney, lung and heart showed no detectable pathological alterations in any of the treatment groups (Figure S9).

Collectively, these data demonstrate that intracochlear delivery of dual AAV-WM04-OTOF vectors is well tolerated across a broad dose range, with minimal auditory impact at supratherapeutic doses and no detectable systemic toxicity, supporting the favorable safety profile of AAV-WM04 for inner ear gene therapy.

### AAV-WM04 enables efficient and safe IHC transduction in NHPs

To assess the translational potential of AAV-WM04, we evaluated the safety, biodistribution, and cochlear transduction efficiency of WM04-CMV-EGFP and WM04-OTOF in cynomolgus macaques. Animals received a single 10 μL intracochlear injection via RWM (Figure S10), and auditory function was monitored longitudinally at 2, 4, and 7 WPI (Figure 5A).

**Figure 5.**
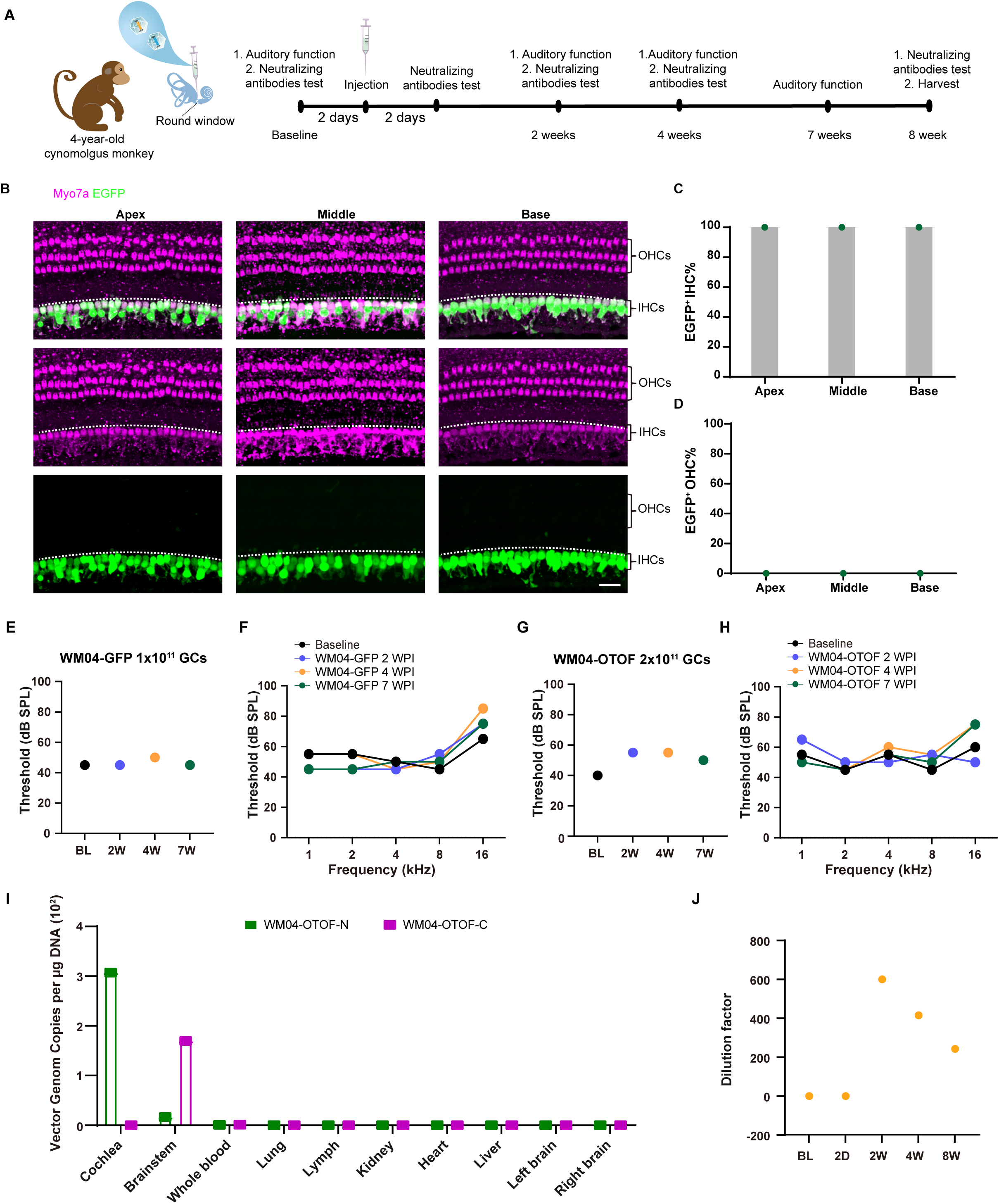
Safety, biodistribution, and inner hair cell specificity of AAV-WM04 in cynomolgus monkeys. **A.** Schematic overview of the safety assessment of dual AAV-WM04-CMV-OTOF vectors administered to cynomolgus monkeys via round window membrane injection. **B.** Representative confocal images of apical, middle, and basal regions of the cochlea following RWM delivery of AAV-WM04-CMV-EGFP (1 × 10¹¹ GCs). Green, EGFP; magenta, MYO7A. Scale bar, 20 μm. **C.** Quantification of EGFP-positive IHCs in the apical, middle, and basal turns of the cynomolgus monkey cochlea corresponding to B. **D.** Quantification of EGFP-positive OHCs in the apical, middle, and basal regions of the cynomolgus monkey cochlea corresponding to B, demonstrating minimal off-target transduction. **E.** Click-evoked ABR thresholds measured at baseline and at multiple time points following RWM injection of AAV-WM04-CMV-EGFP (1 × 10¹¹ GCs). **F.** Tone-burst ABR thresholds measured at baseline and at multiple time points following RWM injection of AAV-WM04-CMV-EGFP (1 × 10¹¹ GCs). **G.** Click-evoked ABR thresholds measured at baseline and at multiple time points following RWM injection of dual AAV-WM04-OTOF vectors (1 × 10¹¹ GCs). **H.** Tone-burst ABR thresholds measured at baseline and at multiple time points following RWM injection of dual AAV-WM04-OTOF vectors (2 × 10¹¹ GCs). **I.** Distribution of OTOF N- and C-terminal sequences in different organs after WM04-OTOF injection. Major tissues were harvested 8 weeks post-injection, and vector DNA copy numbers were quantified per μg of tissue DNA. **J.** Serum neutralizing antibody titers against AAV-WM04 measured at baseline and at indicated time points following injection of dual AAV-WM04-CMV-OTOF in cynomolgus monkeys.

To assess IHC tropism and transduction efficiency, 1 × 10^11^ GCs of WM04-CMV-EGFP were delivered via RWM. At 8 weeks post-injection, temporal bones were harvested and processed following prolonged decalcification. Immunofluorescence analysis revealed robust and exclusive EGFP expression in inner hair cells, achieving 100% IHC transduction throughout the entire cochlear axis, with no detectable expression in supporting cells or outer hair cells (Figure 5B,C). Importantly, no hair cell loss or structural abnormalities were observed (Figure 5D), indicating highly specific and well-tolerated IHC targeting in NHPs, consistent with observations in mice.

Auditory function remained stable following WM04-CMV-EGFP delivery. Click-evoked ABR thresholds were unchanged at 2 and 7 weeks post-injection (45 dB at all time points), consistent with baseline levels (Figure 5E). Frequency-specific ABR thresholds likewise showed no significant differences across baseline, 2, 4, and 7 weeks post-injection (Figure 5F), indicating preserved cochlear function.

To further assess translational safety of the therapeutic construct, 2 × 10¹¹ GCs of dual AAV-WM04-OTOF (1 × 10^11^ GCs each of OTOF-N and OTOF-C) were delivered via RWM. Quantitative PCR analysis demonstrated high levels of OTOF-N and OTOF-C genomes in the cochlea (∼3 × 10^2^ copies per μg DNA), while only low-level OTOF-C genomes were detected in the brainstem (∼1.5 × 10^2^ copies per μg DNA), and OTOF-N was undetectable (Figure 5I). No vector genomes were detected in peripheral organs, including blood, lung, liver, kidney, or heart, indicating limited systemic dissemination.

Functionally, click ABR thresholds showed a transient elevation at 2 weeks post-injection (from 40 dB to 55 dB) but recovered to near-baseline levels by 7 weeks (50 dB) (Figure 5G). Frequency-specific ABR thresholds remained largely unchanged over time, with only a mild, frequency-restricted elevation at 16 kHz, consistent with a non-adverse, dose-limiting observation at supratherapeutic exposure (Figure 5H).

Systemic safety was further evaluated by hematological, biochemical, and inflammatory marker analyses. Although minor fluctuations were observed in individual parameters, none were clinically significant (Table S1; Table S2). Histopathological examination of major organs, including brain, heart, liver, kidney, and intestine, revealed no morphological or pathological abnormalities at 7 weeks post-injection (Figure 5I; Figure S11).

Serum neutralizing antibodies were detected, peaking at 2 weeks post-injection (1:600) and declining to 1:200 by 7 weeks (Figure 5J), consistent with a well-tolerated and self-limited immune response.

Collectively, these data demonstrate that AAV-WM04 mediates highly efficient, specific, and durable IHC transduction in NHPs with a favorable safety and biodistribution profile, strongly supporting its translational potential for clinical inner ear gene therapy.

## Discussion

The successful restoration of hearing through gene therapy offers a transformative therapeutic opportunity for patients with hereditary deafness, particularly those caused by mutations in the *OTOF* gene. A key determinant of clinical translation is the availability of a safe, efficient, and cell type-specific viral vector capable of targeting IHCs, the primary cellular substrate of *OTOF*-related auditory neuropathy. In this study, we established an *in vivo*-directed AAV capsid evolution platform and identified AAV-WM04, a next-generation vector that enables highly efficient and selective transduction of cochlear IHCs across species, resulting in robust and durable hearing restoration in relevant disease models.

Notably, AAV-WM04, identified through *in vivo* screening in the mouse cochlea, retained its high efficiency and specificity for IHCs in NHPs. This cross-species consistency validates the *in vivo-*directed evolution strategy and suggests that the selective pressures applied during screening capture conserved biological determinants of IHC susceptibility rather than rodent-specific features. Given the substantial anatomical and molecular differences between rodent and primate cochleae, the preservation of WM04 tropism underscores the translational relevance of this approach and establishes AAV-WM04 as both a robust IHC-targeted vector and a proof-of-concept for *in vivo* capsid evolution as a broadly applicable platform.

Vector safety represents a central challenge in inner ear gene therapy. Although cell-specific promoters can partially restrict transgene expression, the high viral doses often required for therapeutic efficacy increase the risk of off-target transduction and systemic exposure. In contrast, AAV-WM04 achieves efficient and near-complete IHC transduction at substantially lower doses, thereby expanding the therapeutic window and reducing potential safety liabilities. Consistent with this, no detectable off-target expression was observed in the liver or brain, and auditory function remained intact in control animals, supporting a favorable safety profile.

The inner ear is filled with endolymph and perilymph, creating a fluidic environment that is well suited for AAV perfusion. However, the total volume of the human inner ear is limited (∼282 µL)^32^, which constrains the allowable AAV injection dose. Previous clinical trials have employed injection volumes ranging from 30 to 240 µL via the RWM. Such large infusion volumes inevitably increase intracochlear pressure, often necessitating stapes puncture to facilitate pressure release^11-14^. These additional surgical manipulations, however, may elevate the risk of procedure-related hearing loss. Moreover, due to the limited achievable titers of AAV vectors, reducing the injection volume while maintaining therapeutic efficacy remains technically challenging. Remarkably, less than 10^9^ GC of AAV-WM04 was sufficient to transduce nearly all IHCs in mice, corresponding to an injection volume of approximately 0.1 μL. This low-dose requirement was also associated with reduced neutralizing antibody responses in NHPs, potentially lowering the risk of inflammation and enabling safer clinical application.

Compared with clinically relevant vectors such as AAV1 and Anc80L65, AAV-WM04 exhibited markedly superior IHC transduction efficiency, particularly at low doses. Importantly, this enhanced efficiency translated directly into therapeutic benefit, enabling robust hearing recovery in *Otof* mutant mice using a dual-AAV gene replacement strategy. The large size of the *OTOF* coding sequence necessitates dual-vector approaches, which typically require higher viral loads and raise concerns regarding toxicity and biodistribution. AAV-WM04 supported efficient dual-AAV recombination and functional otoferlin expression at comparatively low doses, mitigating these risks and further underscoring its translational potential.

Hearing restoration mediated by AAV-WM04 was durable, with functional recovery sustained for at least two months following treatment. While recovery at low and mid frequencies was robust and stable, continued optimization of vector design and delivery strategies may further enhance long-term outcomes at the highest frequencies, which remain particularly vulnerable in progressive hearing loss. Notably, AAV-WM04 demonstrated superior efficacy in restoring high-frequency hearing compared with AAV1, a region traditionally difficult to transduce due to anatomical and fluid-dynamic constraints. The strong and uniform cochlear transgene expression achieved by AAV-WM04 provides a mechanistic basis for these improved outcomes.

Several limitations should be acknowledged. First, the duration of long-term follow-up was relatively limited. Extended observation will be required to fully assess the durability and safety of AAV-WM04-mediated hearing restoration. Second, the non-human primate experiments were performed with only one ear per vector, precluding within-animal comparisons and limiting statistical power. Finally, although high-frequency hearing was robustly restored, recovery at low frequencies was comparatively less complete, indicating that further optimization of vector design and delivery strategies may be necessary to achieve uniform functional recovery across the entire cochlear frequency spectrum.

In summary, AAV-WM04 represents a significant advance in the development of safe, efficient, and low-dose gene therapy for *OTOF*-related deafness. Its strong IHC specificity, cross-species efficacy, and favorable safety profile position AAV-WM04 as a promising candidate for future clinical translation. More broadly, this work establishes a generalizable framework for the rational evolution of cell-specific AAV vectors to enable precision treatment of heredity hearing loss.

## Materials and methods

### Animal

All protocols were approved by Shanghai Jiao Tong University Institutional Animal Care and Use Committee (SH9H-2023-T107-1) and followed the 3R principles. Both male and female animals were used and maintained under a 12h light/dark conditions. Wild-type mice, including C57BL/6J and FVB strains, were used in this research. The *Otof* ^Q829X/Q829X^ mice were generated as previously reported. Both male and female animals were randomly assigned to experimental groups. The number of days since birth was counted from postnatal day 0 (P0). Cynomolgus monkey used in this study was male, 5 years old, weighing 3 kg. The animal was housed individually in stainless-steel cages in an AAALAC-accredited facility under a 12-hour light/dark cycle at 23L±L2L°C with 40–70% humidity and received commercial monkey chow and fresh fruit twice daily. All surgical procedures were conducted under general anesthesia with appropriate analgesia. Pre- and post-operative care was provided by trained veterinary staff to minimize animal discomfort.

### AAV capsid library generation

An AAV2 capsid peptide insertion library was generated by introducing randomized 9-mer peptide sequences into the surface-exposed loop between residues N587 and R588 of the AAV2 capsid, a site previously shown to tolerate peptide insertions. The wild-type AAV2 capsid plasmid was linearized via restriction digestion with *BsiWI* (R0553V, NEB) and *SnaBI* (R0130V, NEB), flanking the insertion site. To construct the 9-mer library, a degenerate oligonucleotide encoding nine randomized amino acids (NNK codons) was synthesized (Suzhou GENEWIZ Biotechnology Co., Ltd.). The insertion cassette, containing the random peptide sequence flanked by homology arms corresponding to the AAV2 *cap* gene, was assembled via homologous recombination. Two overlapping DNA fragments encoding the insertion sequence were amplified by PCR using the following primers: Forward 1: 5’-TTACTGACTCGGAGTACCAGCTCCC-3’; Reverse 1: 5’-CATCTGCGGTAGCTGCTTGTCTMNNMNNMNNMNNMNNMNNMNNMNNM NNGTTGCCTCTCTGGAGGTTGGTAGATAC-3’; Forward 2: 5’-AGACAAGCAGCTACCGCAGATG-3’; Reverse 2: 5’-TTAACCCGCCATGCTACTTATCTAC-3’. PCR products were gel-purified using the QIAquick Gel Extraction Kit (28706, QIAGEN). The purified fragments and linearized AAV2 capsid backbone were assembled using homologous recombination with the NovoRec® plus One Step PCR Cloning Kit (NR005-01A, Novoprotein) according to the manufacturer’s instructions. The assembled library constructs were electroporated into *E. coli* MegaX DH10B cells (C640003, Invitrogen) for transformation and amplification.

### AAV production and purification

All AAV vectors were produced in HEK293T cells using polyethyleneimine (PEI)-mediated transfection. For library-scale packaging, capsid library plasmids were co-transfected with adenoviral helper plasmid (pHelper) at a 0.22 μg: 42 μg ratio per 150-mm culture dish. For recombinant AAV production, vectors were generated via a standard triple-plasmid transfection comprising the transgene vector, the rep-cap plasmid, and pHelper. AAV particles were harvested 96 h post-transfection. Both culture supernatants and cell pellets were collected and processed for virus purification. Cell pellets were subjected to repeated freeze-thaw cycles followed by chloroform extraction to release intracellular virions. Supernatants were combined with cell lysates and precipitated overnight at 4 °C using 10% polyethylene glycol (PEG) 8000 and 1.0 M NaCl. The precipitated virus was pelleted by centrifugation and resuspended in PBS supplemented with Benzonase (Merck) to degrade residual DNA. Crude viral suspensions were purified by iodixanol density gradient ultracentrifugation. Gradients were prepared with 15%, 25%, 40%, and 60% iodixanol solutions, and the virus-containing sample was layered atop the gradient and centrifuged at 350,000L×Lg for 90 min at 10 °C. The 40% iodixanol fraction, which contains intact AAV particles, was collected and subjected to buffer exchange and concentration using centrifugal filters (Amicon Ultra, Millipore). Viral genome titers were quantified by SYBR Green–based quantitative PCR (qPCR) targeting the woodchuck hepatitis virus posttranscriptional regulatory element (WPRE). The primers used were: forward, 5′-GTCAGGCAACGTGGCGTGGTGTG-3′ and reverse, 5′-GGCGATGAGTTCCGCCGTGGC-3′.

### Intracochlear injection in adult mice

Posterior semicircular canal (PSCC) or round window membrane (RWM) injection were performed in adult (P30) C57BL/6J or *Otof* ^Q829X/Q829X^ mice, under general anesthesia using xylazine (20 mg/kg, X1251, Sigma Aldrich-Fluka, St. Louis, MO, USA) and Zoletil50 (50 mg/kg, WK001, Virbac S.A.) injected through intraperitoneal injection. Animals were placed in a lateral position, and a postauricular incision was made to expose the otic bulla. For RWM injection, a small fenestration was created in the bulla to access the round window niche. The membrane was gently punctured using a pulled glass micropipette (5-000-1001-X10, DRUMMOND) with tip diameter ∼20–30 μm by vertical needle pulling instrument (PC-10, NARISHIGE). 2 μL of AAV was delivered at the speed of 169 nl/min into the scala tympani over 1–2 minutes by Nanoliter 2000 micromanipulator (WPI). PSCC was performed by a Bonn micro probe (Fine Science Tools, Foster City, CA) and connected it to a Nanoliter 2000 micromanipulator (WPI) by a glass micropipette (WPI, Sarasota, FL) and a fine polyimide tube (inner diameter 0.0039 inches, outer diameter 0.0050 inches, MicroLumen, Tampa, FL). The tip of the tube was inserted into the PSCC, and the hole was sealed by tissue adhesive (3M Vetbond, St. Paul, MN) to deliver AAVs. The total delivery virus volume was 1 μL for each injection at the speed of 169 nL/min. After injection, the tubing was cut and sealed by tissue adhesive. The skin was closed with 7/0 suture (Surgical Suture, B0117B72).

### *In vivo* screening in mice

A total of 1L×L10^10^ GCs of the AAV capsid peptide insertion library were delivered into the cochleae of P30 mice via microinjection through the PSCC. Fourteen days after injection, cochlear tissues were harvested, and viral genomic DNA was extracted using the DNeasy Blood & Tissue Kit (69506, QIAGEN). To identify enriched peptide motifs, the recovered genomic DNA was subjected to next-generation sequencing (NGS). In parallel, cap sequences representing AAV variants capable of transducing the cochlea were PCR-amplified and used to reassemble a secondary AAV library for subsequent rounds of in vivo selection. Three iterative rounds of selection were performed in total. After each round, the frequency of individual variants was quantified by NGS, and enriched capsid sequences were ranked based on their read counts.

### Western blot

HEK293T cells were seeded in six-well plates and maintained in DMEM supplemented with 10% fetal bovine serum (FBS; S711-001S, Lonsera) and 1% penicillin–streptomycin (CB010, CellorLab) at 37 °C in a humidified incubator with 5% COL. The following day, cells were transduced with AAV-WM04-CMV-OTOF-N and AAV-WM04-CMV-OTOF-C at a multiplicity of infection (MOI) of 1L×L10L in DMEM. After a 4-hour incubation, the medium was replaced with high-glucose DMEM supplemented with 10% FBS and antibiotics, and cells were cultured under the same conditions in the absence of mycoplasma and chlamydia contamination. At 48–96 h post-transduction, cells expressing dual AAV-WM04-CMV-OTOF-HA were lysed in RIPA buffer (P0013B, Beyotime) supplemented with protease inhibitors. Lysates were incubated on ice for 30 minutes with intermittent vortexing. Proteins were separated on 4–12% Tris–glycine SDS–PAGE gels (M41210C, GenScript) at 100LV for 90 minutes and transferred to polyvinylidene fluoride (PVDF) membranes (FFP22, Beyotime). Membranes were blocked and probed with an anti-HA-tag primary antibody (1:1,000; 3724S, Cell Signaling Technology), followed by incubation with horseradish peroxidase (HRP)-conjugated anti-rat/mouse IgG secondary antibodies (1:10,000). Protein bands were detected using enhanced chemiluminescence (ECL) and visualized using a chemiluminescence imaging system. Band intensities were quantified with Image Lab software (Bio-Rad).

### Intracochlea injection in Cynomolgus monkeys

RWM injection was performed in cynomolgus monkeys under general anesthesia. Animals were anesthetized with intravenous propofol (5 mg/kg) and maintained at 0.1–0.5 mg/kg/min. Physiological parameters, including respiratory rate, heart rate, arterial blood pressure, and peripheral oxygen saturation, were continuously monitored throughout the procedure. The postauricular region was shaved and sterilized with 10% povidone–iodine. A 35-mm retroauricular semilunar incision was made to expose the mastoid cortex, and meticulous hemostasis was achieved using bipolar electrocautery. Self-retaining retractors were placed, and all subsequent procedures were performed under an operating microscope. Particular care was taken to preserve surrounding anatomical structures, including the external auditory canal, tympanic membrane, facial nerve, and chorda tympani. After identification of the round window niche, the bony overhang was carefully removed to fully expose the round window membrane. Vector delivery was performed using a Nanoliter 2000 micromanipulator (WPI) coupled to a glass micropipette (WPI, Sarasota, FL), at a constant rate of 0.5 μL/min over 20 minutes. A total volume of 10 μL of either AAV-WM04-CMV-EGFP or dual AAV-WM04-CMV-OTOF was injected into the scala tympani via the RWM. Following injection, the RWM was sealed with a small piece of autologous muscle. The mastoid cavity was packed with bone meal and adjacent soft tissue, and the incision was closed in layers. Postoperatively, animals were closely monitored and provided with appropriate analgesia and supportive care to ensure full recovery.

### Immunofluorescence

For cultured cells, samples were fixed in 4% paraformaldehyde (36314ES76, Yeasen) for 2 h at room temperature. For murine cochleae, temporal bones were dissected and fixed in 4% paraformaldehyde for 2 h at room temperature, followed by decalcification in 0.5 M EDTA (ST066, Beyotime) for 1–2 h at room temperature. Cochleae were then dissected into apical, middle, and basal turns. For non-human primate (NHP) cochleae, temporal bones were fixed in 4% paraformaldehyde at room temperature for 2 weeks, followed by decalcification in 10% (w/v) EDTA for 2 months at room temperature. Cochleae were subsequently segmented into apical, middle, and basal regions.

All specimens were permeabilized and blocked in 10% donkey serum with 0.3% Triton X-100 (X10010, Abcone) in 1×PBS (SH30256.FS, HyClone) for 1 h at room temperature. Samples were incubated with primary antibodies overnight at 4 °C. The following primary antibodies were used: anti-Myosin VIIA (25-6790, Proteus Biosciences, 1:1,000), anti-Otoferlin-N (ab53233, Abcam, 1:200), anti-Otoferlin-C (PA552935, Invitrogen, 1:200), anti-HA (11867423001, Roche, 1:50), and anti-Parvalbumin (195004, Synaptic Systems, 1:200). After washing, samples were incubated with the appropriate fluorophore-conjugated secondary antibodies and mounted with VECTASHIELD antifade mounting medium (H-1000-NB, Novios). Fluorescence signals were visualized using a laser-scanning confocal microscope under identical acquisition settings when comparing different conditions.

### Auditory brainstem responses measurement

Auditory brainstem responses were recorded to assess auditory thresholds in mice using a closed-field setup based on the Tucker-Davis Technologies (TDT) System III platform (TDT, Gainesville, FL, USA). All recordings were performed in a sound-attenuated chamber. Animals were anesthetized with intraperitoneal ketamine (100 mg/kg) and xylazine (10 mg/kg), and subdermal needle electrodes were placed at the vertex (active), mastoid region (reference), and dorsal skin of the buttock (ground). ABRs were evoked using tone burst stimuli delivered via a close-field speaker positioned near the external auditory canal. Stimuli were presented at frequencies ranging from 5.66 to 45 kHz. The sound intensity was decreased in 5 dB SPL steps, starting from 90 dB SPL. At each sound level, 512 responses were averaged using alternating stimulus polarity, following automatic artifact rejection. The electrophysiological signals were amplified 10,000-fold, bandpass filtered between 0.1 and 3 kHz, and digitized using the BioSigRZ acquisition software (TDT). A total of 1,024 responses were averaged to generate the final waveform. The auditory threshold was defined as the lowest sound pressure level at which a reproducible and visually detectable ABR waveform could be identified.

### Tissue DNA biodistribution

Following euthanasia, tissue samples (20–25 mg each) were harvested and immediately snap-frozen in liquid nitrogen. Genomic DNA was extracted using a commercial DNA extraction kit (B518251, Sangon Biotech) according to the manufacturer’s instructions. Quantitative PCR (qPCR) was performed to assess the biodistribution of vector DNA across tissues. For the detection of dual AAV constructs, separate primer–probe sets were used to target the N- and C-terminal segments of the OTOF gene. The sequences were as follows: OTOF-N: Forward: 5’-GACGCCATCCACGCTGTTTT-3’; Reverse: 5’-CGAGACTGTCTTGAGGTGGATG-3’; Probe: 5’-VIC652 AGCAAGGCCATGGTGGCGGATC-3’-BHQ1; OTOF-C: Forward: 5’-TGGCTACATGGTCAAAAAGCTC-3’; Reverse: 5’-AGCGTATCCACATAGCGTAAAA-3’; Probe: 5’-FAM-CTTGGGGCATGAGATATCAAGCTTATCG-3’-BHQ1. Reactions were performed on a CFX Real-Time PCR Detection System (Bio-Rad) using standard TaqMan conditions. Vector genome copy numbers were determined by interpolation from a standard curve generated from serial dilutions of plasmid DNA.

### HE Staining

All mouse tissues were fixed in 4% PFA at room temperature for 2 h. Cochlear tissues were similarly decalcified in 10% EDTA prior to dehydration. Samples were dehydrated through a graded ethanol series (30%, 50%, 70%, 80%, 90%, 95%, and 100%), cleared in xylene, embedded in paraffin, and sectioned at 10 μm thickness. Paraffin sections were stained with H&E following standard protocols. Images were captured using a Nikon ECLIPSE Ti inverted microscope.

### AAV neutralizing antibody assay

The titer of anti-AAV neutralizing antibody was determined before and after the administration of EA0010 in all participants. 5×10^4^ HEK-293T cells in 50 μL complete media (DMEM (Gibco), 10% fetal bovine serum (Gibco), 1% penicillin-streptomycin (Sigma-Aldrich)) were seeded per well into a 96-well plate and incubated for 72 h at 37°C. Participant’s blood was collected before (baseline) and at defined time points following the administration of EA0010. Each dilution (3-fold gradient dilution) of the participant’s sera (60 μL) was mixed with 60 μL AAV-EGFP Solution (AAV expressing enhanced green fluorescent protein (EGFP) driven by CMV promotor, Obio Biotechnology) for 1 hour at room temperature with shaking at 600 rpm before adding to the cultured HEK-293T cells. 60 μL of the incubated sample was added into the 96-well plate each well, shaken gently for 1 minute, and then incubated at 37L for 72 ± 4 hours. The green fluorescent signal from expressed EGFP was quantified using a microplate reader (SpectraMax M5) The measurement parameters were set as follows: excitation wavelength, 485 nm; emission wavelength, 525 nm. The gain was adjusted to avoid signal saturation. Wells containing HEK-293T cells without AAV-EGFP infection were used as blank controls. Negative serum of AAV neutralizing antibody mixed with AAV-EGFP were used as negative controls. The relative fluorescence units (RFU) from sample wells were recorded, and the background signal from control wells was subtracted for subsequent analysis. The titer of the neutralizing antibody was defined as the closest dilution of serum that yielded over 50% inhibition of RLU relative to the negative control. The percentage of inhibition was calculated as follows:

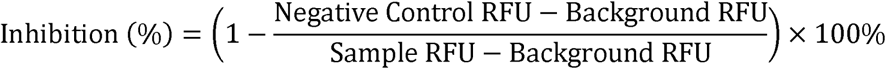

### Rotarod Test

The rotarod apparatus was operated at a constant speed of 8 rpm during the adaptation phase. Mice underwent adaptive training for 30 min per day over three consecutive days. For formal testing, the apparatus was set to an accelerating mode, with the rotational speed increasing linearly from 4 to 40 rpm over a 5-min period. The latency to fall and the rotational speed at the time of falling were recorded. Each mouse was tested in triplicate, and the mean value was used for analysis. A minimum inter-trial interval of 1 hour was maintained to minimize fatigue-related effects. The apparatus was thoroughly cleaned and disinfected between animals to eliminate potential olfactory confounders.

### Pole Test

The pole test was performed using a 1 meter wooden pole inclined at 30° from the horizontal, with the lower end positioned over a cushioned box. Mice were placed head-down on a spherical platform at the top of the pole. The latency was defined as the time from initial contact with the platform to the moment the forepaws reached the ground. Trials were excluded and repeated if mice paused during descent or climbed upward. Each mouse completed two valid trials separated by a 4 hour inter-trial interval, and the average latency was calculated.

### Open Field Test

Mice were individually placed in the center of an open-field arena (50 × 50 × 40 cm) and allowed to explore freely for 30 min. Locomotor activity was continuously recorded using an overhead video camera. Video-tracking software was used to quantify total distance traveled and total time spent in active movement. The arena was cleaned with 75% (v/v) ethanol between trials to prevent cross-contamination.

### Y-Maze Test

The Y-maze consisted of three enclosed arms (38 cm long, 5 cm wide, and 10 cm high) arranged at 120° angles from a central junction. A food reward was placed at the distal end of one designated arm. Mice were introduced into a different arm and allowed to freely explore the maze. The trajectory and latency to reach the food reward were recorded for each trial. The apparatus was cleaned with 75% (v/v) ethanol between trials to remove residual olfactory cues.

### Statistical Analysis

Statistical data were collected and analyzed in GraphPad Prism 9 (GraphPad Software, Inc.). Two-tailed unpaired Student’s *t*-test or one-way ANOVA was used to determine the statistical significance of differences. A value of *P* < 0.05 was considered statistically significant.

## Data and materials availability

All data and materials associated with this study are presented in the paper or in the supplemental information.

## Acknowledgements

We are grateful to the staff members of Bio-imaging core of iHuman Institute, ShanghaiTech University for their technical assistance, and staff members of the Animal Facility at ShanghaiTech University and Kunming Institute of Zoology, Chinese Academy of Sciences for providing the support in animal housing and care.

Hao Wu was supported by Key Project of the National Natural Science Foundation of China (82430037), Natural Science Foundation of Shanghai (24J12800300), Shanghai Key Laboratory of Translational Medicine on Ear and Nose diseases (14DZ2260300).

Yong Tao was supported by the National Natural Science Foundation of China (NSFC82371145, 821220019).

Guisheng Zhong was supported by the National Key Research and Development Program of China (2023YFC3403400) and STI2030-Major Projects (2021ZD0200900), the Science and Technology Commission of Shanghai Municipality (YDZX20223100001002), the Double First-Class Initiative Fund of ShanghaiTech University, and Shanghai Local College Capacity Building Project (22010202700), Shanghai Frontiers Science Center for Biomacromolucules and Precision Medicine and the "Open Competition to Select the Best Candidates" Key Technology Program for Nucleic Acid Drugs of NTICB (Grant No. NCTIB2022HS01009).

## Author Contributions

YT, GZ, and HW conceived and supervised the project and designed and performed the key experiments. YT, CC, ZC, YC, HZ, BY, XH, QY, XM, QZ, and ZD conducted the mouse experiments. BF, CJ, SB, and YS performed data analysis. YT, CC, and ZC drafted the manuscript. All authors critically revised the manuscript for important intellectual content and approved the final version.

## References

1. Yasunaga, S., Grati, M., Cohen-Salmon, M., El-Amraoui, A., Mustapha, M., Salem, N., El-Zir, E., Loiselet, J., and Petit, C. (1999). A mutation in OTOF, encoding otoferlin, a FER-1-like protein, causes DFNB9, a nonsyndromic form of deafness. Nat Genet 21, 363–369. 10.1038/7693.

2. Tang, H., Wang, H., Wang, S., Hu, S.W., Lv, J., Xun, M., Gao, K., Wang, F., Chen, Y., Wang, D., et al. (2023). Hearing of Otof-deficient mice restored by trans-splicing of N- and C-terminal otoferlin. Hum Genet 142, 289–304. 10.1007/s00439-022-02504-2.

3. Sloan-Heggen, C.M., Bierer, A.O., Shearer, A.E., Kolbe, D.L., Nishimura, C.J., Frees, K.L., Ephraim, S.S., Shibata, S.B., Booth, K.T., Campbell, C.A., et al. (2016). Comprehensive genetic testing in the clinical evaluation of 1119 patients with hearing loss. Hum Genet 135, 441–450. 10.1007/s00439-016-1648-8.

4. Rodriguez-Ballesteros, M., Reynoso, R., Olarte, M., Villamar, M., Morera, C., Santarelli, R., Arslan, E., Meda, C., Curet, C., Volter, C., et al. (2008). A multicenter study on the prevalence and spectrum of mutations in the otoferlin gene (OTOF) in subjects with nonsyndromic hearing impairment and auditory neuropathy. Hum Mutat 29, 823–831. 10.1002/humu.20708.

5. Iwasa, Y.I., Nishio, S.Y., Sugaya, A., Kataoka, Y., Kanda, Y., Taniguchi, M., Nagai, K., Naito, Y., Ikezono, T., Horie, R., et al. (2019). OTOF mutation analysis with massively parallel DNA sequencing in 2,265 Japanese sensorineural hearing loss patients. PLoS One 14, e0215932. 10.1371/journal.pone.0215932.

6. Roux, I., Safieddine, S., Nouvian, R., Grati, M., Simmler, M.C., Bahloul, A., Perfettini, I., Le Gall, M., Rostaing, P., Hamard, G., et al. (2006). Otoferlin, defective in a human deafness form, is essential for exocytosis at the auditory ribbon synapse. Cell 127, 277–289. 10.1016/j.cell.2006.08.040.

7. Pangrsic, T., Lasarow, L., Reuter, K., Takago, H., Schwander, M., Riedel, D., Frank, T., Tarantino, L.M., Bailey, J.S., Strenzke, N., et al. (2010). Hearing requires otoferlin-dependent efficient replenishment of synaptic vesicles in hair cells. Nat Neurosci 13, 869–876. 10.1038/nn.2578.

8. Moser, T., and Starr, A. (2016). Auditory neuropathy--neural and synaptic mechanisms. Nat Rev Neurol 12, 135–149. 10.1038/nrneurol.2016.10.

9. Jung, S., Maritzen, T., Wichmann, C., Jing, Z., Neef, A., Revelo, N.H., Al-Moyed, H., Meese, S., Wojcik, S.M., Panou, I., et al. (2015). Disruption of adaptor protein 2mu (AP-2mu) in cochlear hair cells impairs vesicle reloading of synaptic release sites and hearing. EMBO J 34, 2686–2702. 10.15252/embj.201591885.

10. Vincent, P.F., Bouleau, Y., Petit, C., and Dulon, D. (2015). A synaptic F-actin network controls otoferlin-dependent exocytosis in auditory inner hair cells. Elife 4. 10.7554/eLife.10988.

11. Lv, J., Wang, H., Cheng, X., Chen, Y., Wang, D., Zhang, L., Cao, Q., Tang, H., Hu, S., Gao, K., et al. (2024). AAV1-hOTOF gene therapy for autosomal recessive deafness 9: a single-arm trial. Lancet. 10.1016/S0140-6736(23)02874-X.

12. Valayannopoulos, V., Bance, M., Carvalho, D.S., Greinwald, J.H., Jr., Harvey, S.A., Ishiyama, A., Landry, E.C., Lowenheim, H., Lustig, L.R., Manrique, M., et al. (2025). DB-OTO Gene Therapy for Inherited Deafness. N Engl J Med. 10.1056/NEJMoa2400521.

13. Qi, J., Zhang, L., Lu, L., Tan, F., Cheng, C., Lu, Y., Dong, W., Zhou, Y., Fu, X., Jiang, L., et al. (2025). AAV gene therapy for autosomal recessive deafness 9: a single-arm trial. Nat Med 31, 2917–2926. 10.1038/s41591-025-03773-w.

14. Wang, H., Chen, Y., Lv, J., Cheng, X., Cao, Q., Wang, D., Zhang, L., Zhu, B., Shen, M., Xu, C., et al. (2024). Bilateral gene therapy in children with autosomal recessive deafness 9: single-arm trial results. Nat Med 30, 1898–1904. 10.1038/s41591-024-03023-5.

15. Chung, Y., Koehler, S.D., Cancelarich, S., Gibson, T.M., Pregernig, G., Becker, L., Artinian, Q.A., Goodliffe, J.W., Pan, N., Quigley, T.M., et al. (2025). Functional, sustained recovery of hearing in Otoferlin-deficient mice using DB-OTO, a hair-cell-specific AAV-based gene therapy. Mol Ther Methods Clin Dev 33, 101577. 10.1016/j.omtm.2025.101577.

16. Xun, M., Fan, X., Wang, H., Zhao, J., Tang, G., Zhang, W., Hu, S., Zhang, L., Wang, D., Chen, Y., et al. (2025). Comparative analysis of RNA versus protein splicing in dual AAV-mediated gene therapy in a mouse model of DFNB9 deafness. Mol Ther. 10.1016/j.ymthe.2025.10.002.

17. Wang, H., Xun, M., Tang, H., Zhao, J., Hu, S., Zhang, L., Lv, J., Wang, D., Chen, Y., Liu, J., et al. (2024). Hair cell-specific Myo15 promoter-mediated gene therapy rescues hearing in DFNB9 mouse model. Mol Ther Nucleic Acids 35, 102135. 10.1016/j.omtn.2024.102135.

18. Hu, S.W., Lv, J., Wang, Z., Tang, H., Wang, H., Wang, F., Wang, D., Zhang, J., Zhang, L., Cao, Q., et al. (2024). Engineering of the AAV-Compatible Hair Cell-Specific Small-Size Myo15 Promoter for Gene Therapy in the Inner Ear. Research (Wash D C) 7, 0341. 10.34133/research.0341.

19. Qi, J., Zhang, L., Tan, F., Zhang, Y., Zhou, Y., Zhang, Z., Wang, H., Yu, C., Jiang, L., Liu, J., et al. (2024). Preclinical Efficacy And Safety Evaluation of AAV-OTOF in DFNB9 Mouse Model And Nonhuman Primate. Adv Sci (Weinh) 11, e2306201. 10.1002/advs.202306201.

20. Rankovic, V., Vogl, C., Dorje, N.M., Bahader, I., Duque-Afonso, C.J., Thirumalai, A., Weber, T., Kusch, K., Strenzke, N., and Moser, T. (2020). Overloaded Adeno-Associated Virus as a Novel Gene Therapeutic Tool for Otoferlin-Related Deafness. Front Mol Neurosci 13, 600051. 10.3389/fnmol.2020.600051.

21. Tertrais, M., Bouleau, Y., Emptoz, A., Belleudy, S., Sutton, R.B., Petit, C., Safieddine, S., and Dulon, D. (2019). Viral Transfer of Mini-Otoferlins Partially Restores the Fast Component of Exocytosis and Uncovers Ultrafast Endocytosis in Auditory Hair Cells of Otoferlin Knock-Out Mice. J Neurosci 39, 3394–3411. 10.1523/JNEUROSCI.1550-18.2018.

22. Akil, O., Dyka, F., Calvet, C., Emptoz, A., Lahlou, G., Nouaille, S., Boutet de Monvel, J., Hardelin, J.P., Hauswirth, W.W., Avan, P., et al. (2019). Dual AAV-mediated gene therapy restores hearing in a DFNB9 mouse model. Proc Natl Acad Sci U S A 116, 4496–4501. 10.1073/pnas.1817537116.

23. Al-Moyed, H., Cepeda, A.P., Jung, S., Moser, T., Kugler, S., and Reisinger, E. (2019). A dual-AAV approach restores fast exocytosis and partially rescues auditory function in deaf otoferlin knock-out mice. EMBO Mol Med 11. 10.15252/emmm.201809396.

24. Zhang, L., Wang, H., Xun, M., Tang, H., Wang, J., Lv, J., Zhu, B., Chen, Y., Wang, D., Hu, S., et al. (2023). Preclinical evaluation of the efficacy and safety of AAV1-hOTOF in mice and nonhuman primates. Mol Ther Methods Clin Dev 31, 101154. 10.1016/j.omtm.2023.101154.

25. Pupo, A., Fernandez, A., Low, S.H., Francois, A., Suarez-Amaran, L., and Samulski, R.J. (2022). AAV vectors: The Rubik’s cube of human gene therapy. Mol Ther 30, 3515–3541. 10.1016/j.ymthe.2022.09.015.

26. Andres-Mateos, E., Landegger, L.D., Unzu, C., Phillips, J., Lin, B.M., Dewyer, N.A., Sanmiguel, J., Nicolaou, F., Valero, M.D., Bourdeu, K.I., et al. (2022). Choice of vector and surgical approach enables efficient cochlear gene transfer in nonhuman primate. Nat Commun 13, 1359. 10.1038/s41467-022-28969-3.

27. Landegger, L.D., Pan, B., Askew, C., Wassmer, S.J., Gluck, S.D., Galvin, A., Taylor, R., Forge, A., Stankovic, K.M., Holt, J.R., et al. (2017). A synthetic AAV vector enables safe and efficient gene transfer to the mammalian inner ear. Nat Biotechnol 35, 280–284. 10.1038/nbt.3781.

28. Tao, Y., Huang, M., Shu, Y., Ruprecht, A., Wang, H., Tang, Y., Vandenberghe, L.H., Wang, Q., Gao, G., Kong, W.J., et al. (2018). Delivery of Adeno-Associated Virus Vectors in Adult Mammalian Inner-Ear Cell Subtypes Without Auditory Dysfunction. Hum Gene Ther 29, 492–506. 10.1089/hum.2017.120.

29. Shu, Y., Tao, Y., Wang, Z., Tang, Y., Li, H., Dai, P., Gao, G., and Chen, Z.Y. (2016). Identification of Adeno-Associated Viral Vectors That Target Neonatal and Adult Mammalian Inner Ear Cell Subtypes. Hum Gene Ther 27, 687–699. 10.1089/hum.2016.053.

30. Chan, K.Y., Jang, M.J., Yoo, B.B., Greenbaum, A., Ravi, N., Wu, W.L., Sanchez-Guardado, L., Lois, C., Mazmanian, S.K., Deverman, B.E., et al. (2017). Engineered AAVs for efficient noninvasive gene delivery to the central and peripheral nervous systems. Nat Neurosci 20, 1172–1179. 10.1038/nn.4593.

31. Xue, Y., Tao, Y., Wang, X., Wang, X., Shu, Y., Liu, Y., Kang, W., Chen, S., Cheng, Z., Yan, B., et al. (2023). RNA base editing therapy cures hearing loss induced by OTOF gene mutation. Mol Ther 31, 3520–3530. 10.1016/j.ymthe.2023.10.019.

32. Inui, H., Kitahara, T., Ito, T., and Sakamoto, T. (2021). Magnetic Resonance 3D Measurement of the Endolymphatic Space in 100 Control Human Subjects. J Int Adv Otol 17, 536–540. 10.5152/iao.2021.21317.

