## supplementary for "*In vivo*-directed evolution identifies AAV-WM04 as a next-generation vector for potent and durable hearing restoration in DFNB9"

#### Figure S1

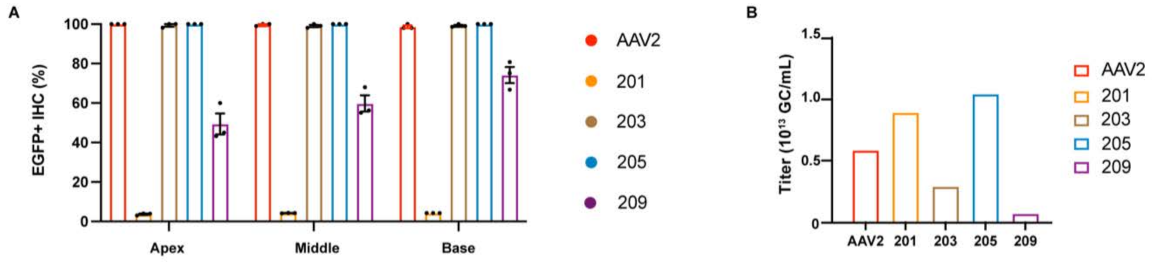


**Figure S1. Transduction efficiency and packaging capacity of AAV2 and candidate AAV vectors.**

**A.** Transduction efficiency of AAV2 and AAV candidate vectors (201, 203, 205, and 209), quantified as the percentage of EGFP-positive IHCs in the apical, middle, and basal turns of the cochlea.

**B.** Maximum viral titers achievable for each AAV vector.

#### Figure S2

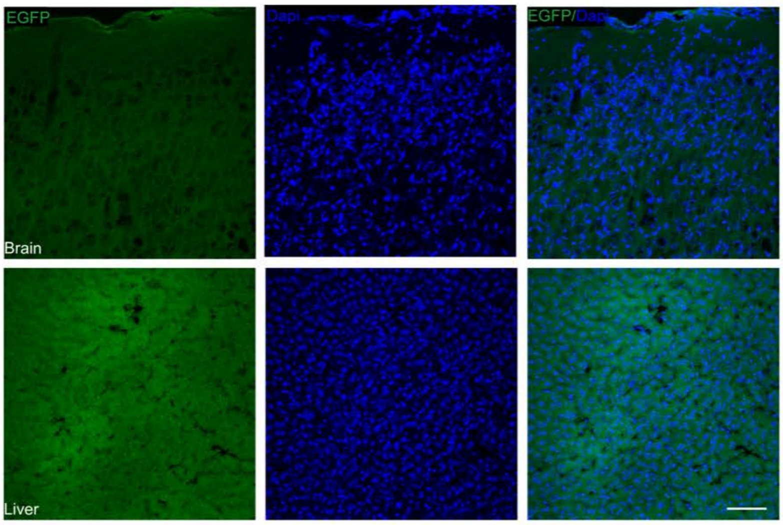


**Figure S2. Absence of EGFP expression in the brain and liver of adult WT mice after AAV-WM04-CMV-EGFP injection.**
Representative confocal images of the brain and liver from adult WT mice at 14 days post-injection with $2 \times 10^9$ GCs AAV-WM04-CMV-EGFP. Blue, DAPI; green, EGFP. Scale bar, 20 µm.

### Figure S3

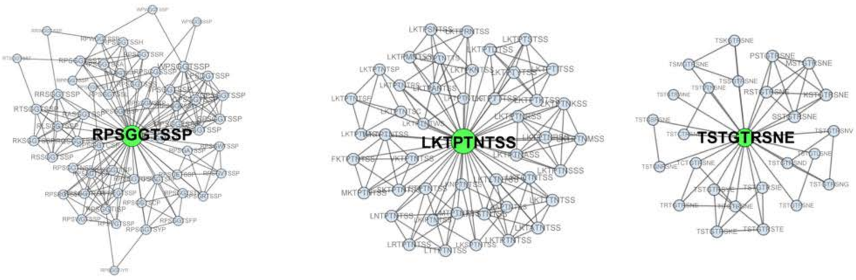


**Figure S3 Data analysis of three rounds screening in adult mice.**
Clustering analysis of 3 typical families (201, 203 and 209) from positively enriched variants across all three rounds.

**Figure S4**

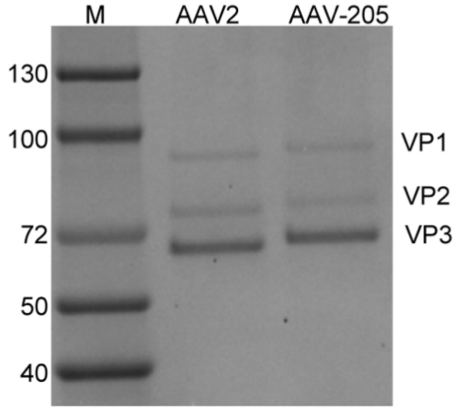


**Figure S4. Purity analysis of AAV2 and AAV-WM04 by SDS-PAGE.** M means Marker.

#### Figure S5

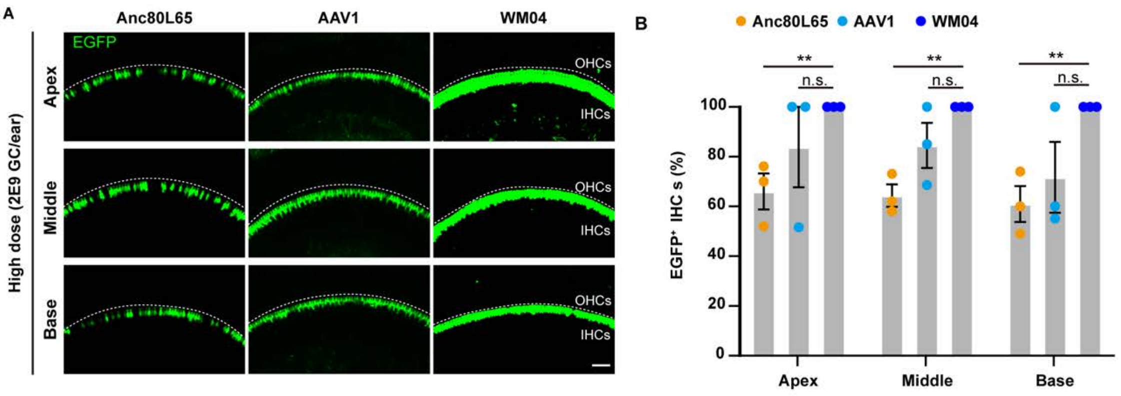


**Figure S5. AAV-WM04-CMV-EGFP efficiently transduces inner hair cells in P30 mice. A.** Representative confocal images of cochlear inner hair cells from adult wild-type mice at 14 days post-injection with $2 \times 10^9$ GCs of AAV-WM04-CMV-EGFP, Anc80L65-CMV-EGFP, or AAV1-CMV-EGFP. Green,EGFP; IHCs, inner hair cells; OHCs, outer hair cells. Scale bar, 20 μm. **B.** Percentage of EGFP-positive IHCs corresponding to A. Data are presented as mean ± SEM; n = 3 mice per group. Statistical significance was determined by Student's *t*-test. n.s., not significant.

#### Figure S6

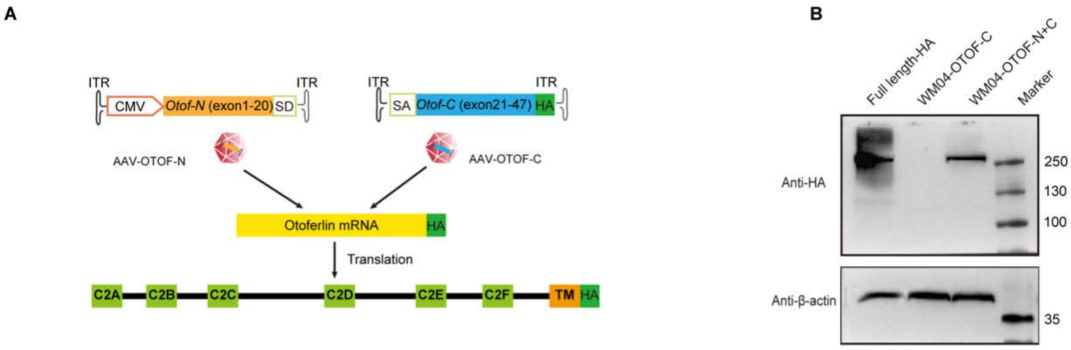


**Figure S6. Dual AAV-WM04-CMV-*OTOF* induced the full-length recombination and expression of OTOF. A.** Schematic representation of the recombinant AAV-vector pair used in this study. Human *Otoferlin* isoform V was split at exon 20/21. The recombinant AAV-*OTOF*-N and AAV-*OTOF*-C vectors contain the 5′ and 3′ parts of the *OTOF* cDNA. In addition, the splicing donor (SD) and splicing acceptor (SA) elements in the vectors were included for RNA splicing. HA tag was added in the C-terminal human *Otoferlin* for detection. Abbreviations: ITR, inverted terminal repeats; CMV, cytomegalovirus immediate early promoter; C2, C2 domain; TM, transmembrane domain. B**.** Full-length *Otoferlin* expression was detected by western blot after 48h transduction with 1:1 ratio of AAV-WM04-CMV-OTOF-N and AAV-WM04-CMV-OTOF-C-HA in HEK293T cells. $\beta$-actin was used as the reference gene.

### Figure S7

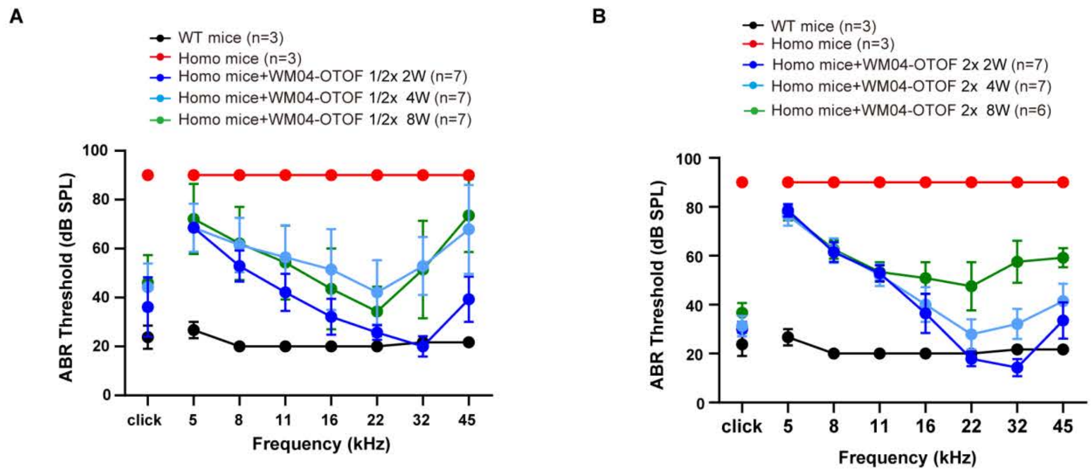


**Figure S7. WM04-*OTOF* injection restored auditory function in P30 *Otof* $^{Q829X/Q829X}$ mice.**

**A.** Click and tone-burst ABR thresholds of P30 wild-type mice (black, n = 3), untreated homozygous mice (red, n = 4) and homozygous mice treated with a low dose of WM04-OTOF (2.5 × $10^9$ GCs, 1/2×) at 2 weeks (blue, n = 7), 4 weeks (light blue, n = 7), and 8 weeks (green, n = 7) post-injection. Data are shown as mean ± SEM. Statistical significance was assessed by two-way ANOVA compared with untreated homozygous mice.

**B.** Click and tone-burst ABR thresholds of P30 wild-type mice (black, n = 3), untreated homozygous mice (red, n = 4) and homozygous mice treated with a low dose of WM04-OTOF (1 × $10^{10}$ GCs, 2×) at 2 weeks (blue, n = 7), 4 weeks (light blue, n = 7), and 8 weeks (green, n = 6) post-injection. Data are shown as mean ± SEM. Statistical significance was assessed by two-way ANOVA compared with untreated homozygous mice.

#### Figure S8

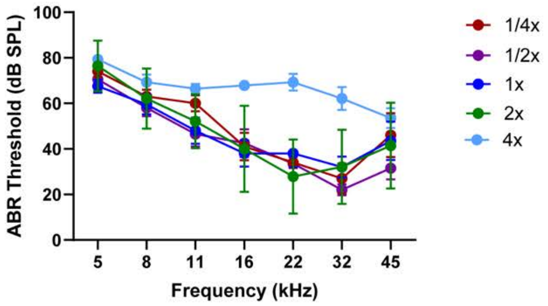


**Figure S8. Dose–response analysis of tone-burst ABR thresholds**
Tone-burst ABR thresholds at 4 weeks post-injection in homozygous mice treated with WM04-OTOF at $1.25 \times 10^9$ (1/4×, n = 5), $2.5 \times 10^9$ (1/2×, n = 7), $5 \times 10^9$ (1×, n = 8), $1 \times 10^{10}$ (2×, n = 6), and $2 \times 10^{10}$ GCs (4×, n = 5). Data are shown as mean ± SEM.

#### Figure S9

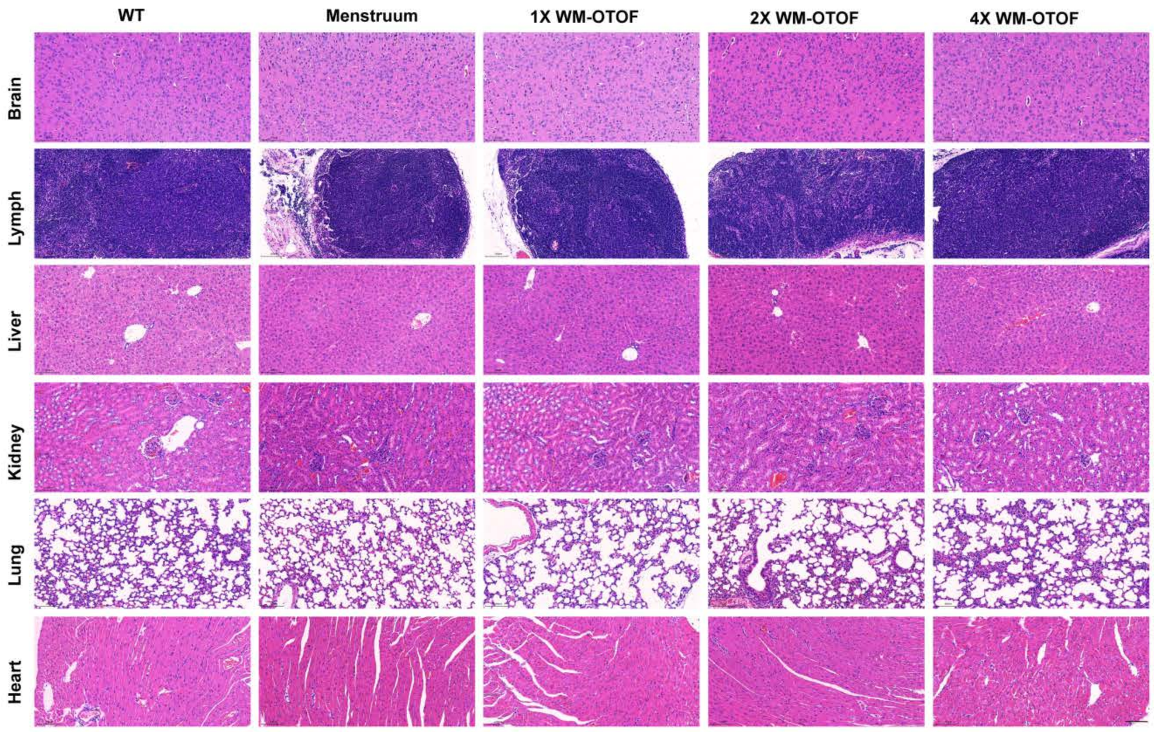


**Figure S9. RWM delivery of WM04-OTOF is well tolerated in wild-type mice without detectable adverse effects.**
Representative H&E-stained sections of major tissues from WT mice following RWM administration of WM04-OTOF. No overt histopathological abnormalities were observed. Scale bar, 100 µm.

### Figure S10

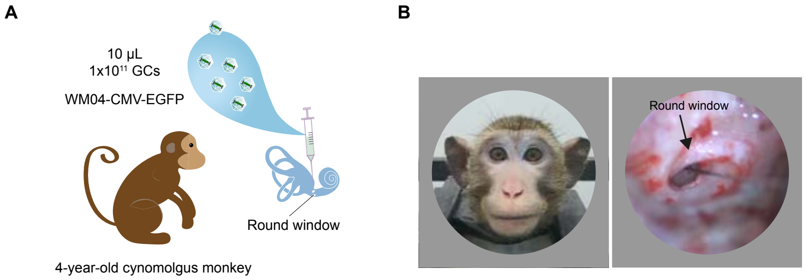


**Figure S10. Efficient transduction of the cynomolgus monkey cochlea by AAV-CMV-WM04-eGFP following RWM delivery.**

A. Schematic overview of the experimental design for AAV-WM04-CMV-EGFP administration in cynomolgus monkeys. B. Representative images showing the cynomolgus monkey and surgical exposure of RWM for AAV delivery.

**Figure S11**

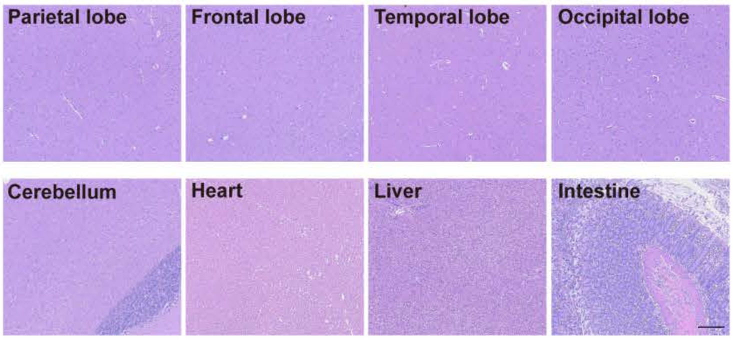


**Figure S11. RWM delivery of WM04-OTOF in cynomolgus monkeys was well tolerated, with no detectable adverse effects.**

H&E-staining of major organs was performed 8 weeks after WM04-OTOF injection, revealing no detectable histopathological abnormalities. Scale bar: 200 μm.

**Table S1. Routine blood test in cynomolgus monkey**

| Time post-injection | Baseline | 2 days | 2 weeks | 4weeks | 8weeks |
|---|---|---|---|---|---|
| White blood cells ($10^9$/L) | 13.5 | 16.5 | 8.2 | 14.9 | 20 |
| Red blood cells ($10^{12}$/L) | 5.57 | 5.98 | 4.35 | 5.38 | 5.97 |
| Hemoglobin (g/L) | 138 | 145 | 125 | 131 | 148 |
| Packed cell volume (%) | 41.3 | 46 | 43.7 | 39.8 | 45.7 |
| Mean corpuscular volume(fl) | 74 | 77 | 100 | 74 | 77 |
| Mean corpuscular hemoglobin (pg.) | 24.7 | 24.2 | 28.7 | 24.4 | 24.8 |
| Mean corpuscular hemoglobin concentration (g/L) | 334 | 315 | 286 | 329 | 324 |
| Red blood cell distribution (%) | 12.1 | 12.5 | 19.3 | 12.6 | 13.4 |
| Platelet ($10^9$/L) | 213 | 356 | 650 | 412 | 377 |
| Mean platelet volume(fl) | 13.8 | 15 | 12.6 | 12.9 | 15.5 |
| Plateletcrit (%) | 0.294 | 0.534 | 0.819 | 0.531 | 0.584 |
| platelet distribution (%) | 16.2 | 15.7 | 16.5 | 16 | 15.5 |
| lymphocyte (%) | 68.2 | 46.9 | 2.5 | 60.2 | 73.7 |
| Lymphocyte ($10^9$/L) | 9.21 | 7.72 | 0.21 | 8.96 | 14.71 |
| Monocyte (%) | 5 | 7.9 | 5.6 | 3.9 | 4.8 |
| Monocyte (($10^9$/L) | 0.68 | 1.3 | 0.46 | 0.58 | 0.96 |
| Neutrophils (%) | 25.2 | 44 | 66.8 | 35 | 19.9 |
| Neutrophils ($10^9$/L) | 3.41 | 7.25 | 5.49 | 5.2 | 3.97 |
| Eosinophil (%) | 1.6 | 0.3 | 0.6 | 0.9 | 1.1 |
| Eosinophil ($10^9$/L) | 0.22 | 0.05 | 0.05 | 0.13 | 0.22 |
| Basophil (%) | 0 | 0.9 | 24.5 | 0 | 0.5 |
| Basophil ($10^9$/L) | 0 | 0.15 | 2.01 | 0 | 0.1 |

**Table S2. Blood biochemistry in cynomolgus monkey**

| Time post-injection | Baseline | 2 days | 2 weeks | 4weeks | 8weeks |
|---|---|---|---|---|---|
| Total protein (g/L) | 66.4 | 83.9 | 66.5 | 65.2 | 72 |
| C-reactive protein (mg/L) | 1 | 21.1 | 4.3 | 7.7 | 4.3 |
| Lactate dehydrogenase (U/L) | 987 | 797 | 1199 | 1518 | 326 |
| Creatine kinase (U/L) | 384 | 1061 | 167 | 243 | 192 |
| Alkaline phosphatase (U/L) | 635 | 699 | 536 | 539 | 502 |
| Triglyceride (mmol/L) | 0.6 | 0.33 | 0.54 | 0.79 | 0.82 |
| Serum total cholesterol (mmol/L) | 2.58 | 2.5 | 2.23 | 1.96 | 2.73 |
| Total bilirubin (μmol/L) | 1.3 | 2.9 | 1.6 | 2.1 | 0.9 |
| Albumin (g/L) | 35.5 | 40.4 | 31.4 | 33.3 | 37.7 |
| Alanine aminotransferase (U/L) | 32 | 78 | 87 | 46 | 25 |
| Aspartate aminotransferase (U/L) | 63 | 111 | 84 | 85 | 36 |
| Crea (μmol/L) | 77 | 75 | 54 | 98 | 77 |
| Urea (mmol/L) | 6.32 | 5.58 | 5.81 | 2.17 | 4.56 |

#### Table S3 Distribution of OTOF N- and C-terminal sequences

| | Cochlea | Brain-stem | Whole blood | Lung | Lymph | Kidney | Heart | Liver | Left brain | Right brain |
|---|---|---|---|---|---|---|---|---|---|---|
| OTOF-N | 307 | 16.5 | 0.9 | 0.3 | 0.2 | 0.2 | 0.1 | 0.1 | 0.1 | 0.1 |
| OTOF-C | BDL | 169.7 | 1 | BDL | BDL | BDL | BDL | 0.1 | 0.1 | BDL |

Unit: DNA copy numbers/μg of tissue DNA
BDL: below the detection limit
